# Trajectory reconstruction identifies dysregulation of perinatal maturation programs in pluripotent stem cell-derived cardiomyocytes

**DOI:** 10.1101/2021.01.31.428969

**Authors:** Suraj Kannan, Matthew Miyamoto, Brian L. Lin, Chulan Kwon

## Abstract

A primary limitation in the clinical application of pluripotent stem cell derived cardiomyocytes (PSC-CMs) is the failure of these cells to achieve full functional maturity. *In vivo*, cardiomyocytes undergo numerous adaptive changes during perinatal maturation. By contrast, PSC-CMs fail to fully undergo these developmental processes, instead remaining arrested at an embryonic stage of maturation. To date, however, the precise mechanisms by which directed differentiation differs from endogenous development, leading to consequent PSC-CM maturation arrest, are unknown. The advent of single cell RNA-sequencing (scRNA-seq) has offered great opportunities for studying CM maturation at single cell resolution. However, perinatal cardiac scRNA-seq has been limited owing to technical difficulties in the isolation of single CMs. Here, we used our previously developed large particle fluorescence-activated cell sorting approach to generate an scRNA-seq reference of mouse *in vivo* CM maturation with extensive sampling of perinatal time periods. We subsequently generated isogenic embryonic stem cells and created an *in vitro* scRNA-seq reference of PSC-CM directed differentiation. Through trajectory reconstruction methods, we identified a perinatal maturation program in endogenous CMs that is poorly recapitulated *in vitro*. By comparison of our trajectories with previously published human datasets, we identified a network of nine transcription factors (TFs) whose targets are consistently dysregulated in PSC-CMs across species. Notably, we demonstrated that these TFs are only partially activated in common *ex vivo* approaches to engineer PSC-CM maturation. Our study represents the first direct comparison of CM maturation *in vivo* and *in vitro* at the single cell level, and can be leveraged towards improving the clinical viability of PSC-CMs.

**Significance Statement:** There is a significant clinical need to generate mature cardiomyocytes from pluripotent stem cells. However, to date, most differentiation protocols yield phenotypically immature cardiomyocytes. The mechanisms underlying this poor maturation state are unknown. Here, we used single cell RNA-sequencing to compare cardiomyocyte maturation pathways in endogenous and pluripotent stem cell-derived cardiomyocytes. We found that *in vitro*, cardiomyocytes fail to undergo critical perinatal gene expression changes necessary for complete maturation. We found that key transcription factors regulating these changes are poorly expressed *in vitro.* Our study provides a better understanding of cardiomyocyte maturation both *in vivo* and *in vitro*, and may lead to improved approaches for engineering mature cardiomyocytes from stem cells.

---

Pluripotent stem cell (PSC)-derived cardiomyocytes (CMs) offer a powerful solution to numerous challenges in clinical cardiology, with applications including regenerative medicine, drug screening, and disease modeling (1, 2). However, the inability of PSC-CMs to mature to an adult-like phenotype has precluded their effective biomedical use (3, 4). A number of transcriptomic, proteomic, structural, and functional measurements have indicated that PSC-CMs more closely resemble fetal or embryonic CMs rather than their mature adult counterparts (3, 5–7). While this phenomenon is often termed a “maturation arrest,” there is little mechanistic understanding of why PSC-CMs fail to recapitulate the adult phenotype. It is known that differentiating PSC-CMs faithfully undergo cascading gene expression changes associated with gastrulation, mesoderm induction, and cardiomyogenesis (8, 9). However, it is unclear to what degree they initiate a maturation-like program *in vitro*, or when and how this program is disrupted. This is further complicated by the lack of knowledge of the initiation, regulation, and dynamics of CM maturation *in vivo* (4). To address this, our previous work aimed to identify a transcriptional landscape for CM maturation over *in vivo* development (10). We subsequently compared the establishment of gene regulatory networks (GRNs) between PSC-CMs and endogenous CMs. However, this work was done using bulk samples, while CM maturation occurs heterogeneously across time both *in vivo* and *in vitro* (5, 11), suggesting a need for further data.

To address the limited maturation of PSC-CMs, a number of groups have developed *ex vivo* perturbation protocols to improve PSC-CM maturation. These approaches have included cytokine, growth factor, and hormone cocktails, co-culture with other cells, induction of physical stimuli (e.g. mechanical stretch, electrical stimulation), and construction of biomaterial-based three dimensional tissues (3, 12–16). Ostensibly, the goal of these perturbations is to replicate fundamental aspects of the native cardiac milieu to engineer maturation. However, assessment of the maturation of these perturbed tissues is often done through *ad hoc* phenotypic measurements. Additionally, there has been no systematic comparison of whether these perturbations activate maturation pathways analogous to endogenous development. Thus, it is unclear to what degree any individual perturbation is truly biomimetic. Other groups have shown that, despite their immature phenotype, PSC-CMs may already be capable of modeling specific (albeit limited) aspects of cardiac biology. This approach has led to breakthrough applications in drug screening and disease modeling (17, 18). A deeper biological understanding of CM maturation processes, however, can significantly expand the possibilities for future clinical application of PSC-CMs.

In this study, we used single cell RNA-sequencing (scRNA-seq) to directly compare maturation processes between endogenous and PSC-derived CMs. Through use of our previously established large-particle fluorescence-activated cell sorting (LP-FACS) protocol (19), we established a high quality reference of mouse CM maturation, in particular sampling previously understudied perinatal timepoints. We subsequently generated embryonic stem cell (ESC) lines from the same strain used for the *in vivo* reference, and differentiated these to PSC-CMs to produce an *in vitro* maturation reference. We found that endogenous CMs undergo a perinatal maturation program between postnatal day (p)8-p15 that is poorly recapitulated *in vitro*. By cross-referencing with published human PSC-CM datasets, we identified a network of nine transcription factors (TFs) whose targets are consistently dysregulated in PSC-CMs across species. Through further meta-analysis of published perturbation RNA-seq datasets, we found that *ex vivo* perturbations can partially activate some of the key maturation-related TFs, but no method completely activates them all. Our study is the first to provide systematic single cell comparison of maturation programs between *in vivo* and *in vitro* CMs, and may open future avenues for generating fully mature PSC-CMs.

## Results

### Trajectory reconstruction identifies perinatal window in endogenous CM maturation

Our first goal was to develop an improved understanding of gene expression changes in CM maturation during *in vivo* development. scRNA-seq of CM maturation, in particular postnatal maturation, has been previously limited due to difficulty in isolating large, fragile CMs (20). Recently, however, we established LP-FACS as a viable approach to generate high-quality CM scRNA-seq libraries (19), and used this technique to produce a reference dataset of CM maturation (5). In this study, we expanded our reference to a total of ~ 1600 left ventricular free wall CMs encompassing 15 timepoints ranging from embryonic day (e)14 to p84 (**Supplementary Note 1**). We specifically isolated CMs through use of Myh6-Cre; mTmG (*α*MHC x mTmG) mice, in which cells expressing cardiac-specific myosin heavy chain are readily separated by GFP expression (**Figure 1A**). We further validated CM identity of sequenced libraries through use of the SingleCellNet algorithm (21), which computationally classifies single cells by gene expression (**Figure S1A**). CMs were sequenced to a median depth of ~21,000 unique molecular identifiers (UMIs) per cell and ~4000 genes per cell. This high-depth sequencing is particularly important for analyzing CMs undergoing maturation because of their gradual transcriptomic changes. Our dataset was designed to encompass the full temporal range of CM maturation while particularly sampling perinatal timepoints that may be critical to the maturation process (4, 22).

**Fig. 1.**
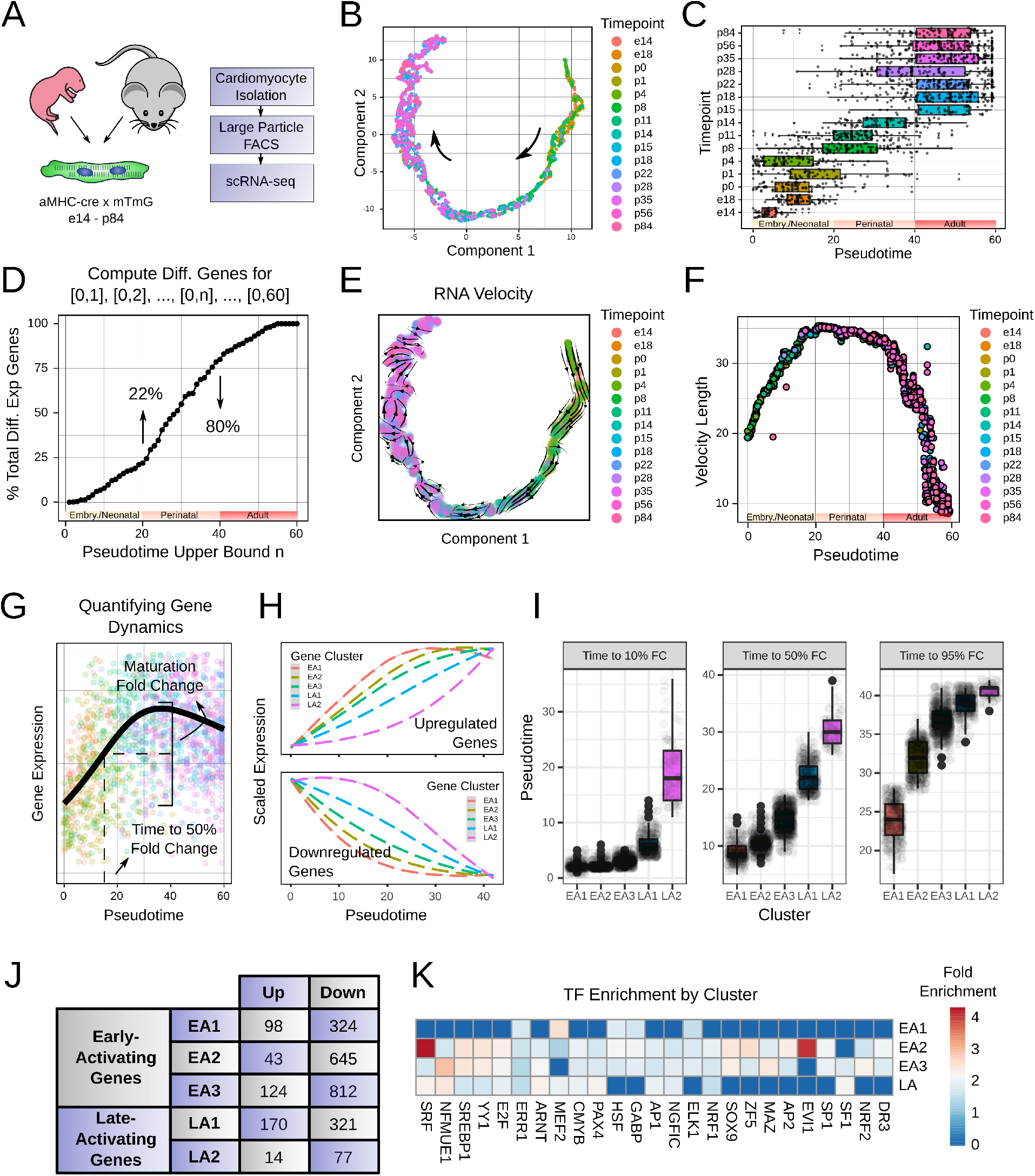
scRNA-seq data reconstructs a developmental trajectory of in vivo CM maturation. **A.** Schematic of the experimental workflow for generating *in vivo* CM libraries. **B.** Trajectory inferred by Monocle 3, labeled by timepoint. Arrow indicates direction of pseudotime. **C.** Pseudotime scores per timepoint for the inferred trajectory. We designated pseudotime intervals as embryonic/neonatal, perinatal, and adult respectively. **D.** Percent of the total differentially expressed genes that become differentially expressed within a pseudotime interval. We subdivided pseudotime into binned subsets of cells and identified differentially expressed genes for each subset. **E.** RNA velocity stream plot projected onto inferred trajectory, labeled by timepoint. **F.** RNA velocity length across Monocle 3-inferred pseudotimes. **G.** Schematic of the workflow for computing gene dynamics. We defined the maturation FC as the maximum FC in the pseudotime interval [0, 42], and computed the time to achieve a designated percentage of this FC. **H.** Average gene dynamics for each gene cluster, split into upregulated and downregulated genes for each cluster. Clusters were determined based on gene dynamics parameters. **I.** Gene dynamics parameters (time to 10%, 50%, and 95% FC) for each cluster. EA = early activation, LA = late activation, with groupings based on the time to 10% FC. **J.** Number of upregulated and downregulated genes in each identified cluster. **K.** TFs whose downstream targets are enriched in each gene cluster. As LA2 has relatively few genes, we combined LA1 and LA2 and performed enrichment.

To understand the gene expression changes over CM maturation at the single cell level, we performed trajectory reconstruction using Monocle 3 (23) (**Figure 1B, Supplementary Note 2**). Monocle 3 recovered a unidirectional, non-branching trajectory, matching earlier reports (5, 11, 24). Trajectory reconstruction enables calculation of “pseudotime,” a metric of progression along an inferred biological process. We plotted the Monocle 3-computed pseudotimes at each biological timepoint, which validated that pseudotime progressively increased over biological time (**Figure 1C**). Notably, however, at each biological timepoint, there was significant heterogeneity of pseudotime, indicating that maturation proceeds asynchronously at single cell level. Additionally, we observed several interesting transition points. From e14 to e18, there was a jump in pseudotime marking the initiation of maturation. Between e18 and p4 (corresponding to pseudotime ∈ [5,20]), individual cells proceeded through a late embryonic/neonatal phase of maturation, but the distribution of pseudotimes was similar at each biological timepoint. Subsequently, there was a large jump in pseudotime between p8 and p14 (corresponding to pseuodotime ∈ (20,40]), suggesting a perinatal maturation process. Finally, after p15 (corresponding to pseudotime ∈ (40, 60]), cells converged to a relatively mature phenotype, though there was still notable heterogeneity even in these late stages. Based on these results, we labeled the pseudotime intervals [0, 20], (20, 40], and (40, 60] as embryonic/neonatal, perinatal, and adult respectively.

Using Monocle 3, we identified 3015 genes with differential expression over pseudotime. Over 80% of these genes were downregulated over pseudotime, which supports our previous assertion that CM maturation involves significant pruning of unneeded gene modules (5). As expected, upregulated genes corresponded to crucial aspects of mature CM biology, including sarcomeric, calcium handling, and fatty acid metabolism, while downregulated genes were enriched for cell cycle and transcription/translation processes (**Figure S1B**). We next aimed to identify when maturation genes become differentially expressed along pseudotime. To do this, we produced sequential subsets of our inferred trajectory by binning cells with pseudotime ∈ [0, 1], pseudotime ∈ [0, 2] etc. up to a bin of cells with pseudotime ∈ [0, 60] (e.g. the entire trajectory). We then computed differential genes for each subset, to identify what percentage of the total 3015 differentially expressed genes become differentially expressed within each progressive subset. 22% of genes become differentially expressed by pseudotime 20 while 80% become differentially expressed by pseudotime 40 (**Figure 1D**). Thus, 58% of genes become differentially expressed within the perinatal period, further supporting this period as critical for CM maturation.

As a complementary approach to trajectory reconstruction, we investigated transcriptional dynamics by computing RNA velocity using the scVelo package (25). RNA velocity uses the ratio of spliced and unspliced messenger RNA reads in scRNA-seq data to determine the rate of gene expression changes (26). These computed velocities can be aggregated and projected onto developmental trajectories to study cell differentiation dynamics. Here, we computed velocities for our identified differentially expressed genes, and embedded these velocities on our Monocle 3-inferred trajectory (**Figure 1E**). The RNA velocity field pointed along the maturation trajectory in the embryonic and perinatal phases before subsequently becoming more incoherent. The loss of coherence of the velocity field in later timepoints likely indicates that these timepoints at dynamic equilibrium with no clear transcriptional directionality, suggesting completion of maturation. These results are further supported by the ratios of spliced to unspliced RNAs, which steadily decrease over biological time until becoming stable around p15 (**Figure S1C**). The rate of differentiation can be quantified by the length of the velocity vector. We found that the velocity length progressively increased starting at pseudotime 0, reaching a maximum for pseudotime ∈ [20, 40], before subsequently decreasing (**Figure 1F**). These results indicate that endogenously, CM maturation proceeds with highest velocity in the perinatal phase.

Taken together, our results support the existence of a perinatal window for CM maturation. Based on this, we limited our subsequent analysis to the perinatal period by selecting cells with pseudotime ∈ [0, 42]. We selected 42 as our upper pseudotime threshold to effectively capture our identified perinatal window ((20, 40]) plus a small margin of error. During this period, 2628 genes become differentially expressed, which we subsequently refer to as maturation genes.

### Maturation genes form early-activating and late-activating clusters with unique upstream regulation

We next sought to determine and isolate groups of genes that share temporal expression patterns from embryonic to perinatal stages. Though some previous methods have been developed for quantifying gene dynamics along pseudotime trajectories (27), we instead developed a flexible method using built-in features of Monocle 3. For each gene, Monocle 3 fits a smooth spline curve (**Figure 1G**). We defined the maturation fold change (FC) as the maximum fold change that happens over this spline curve in our predetermined pseudotime interval of [0, 42]. We then identified the time to x% fold change as the pseudotime at which x% of the maturation fold change is achieved. The x% used can be calibrated to study different aspects of individual gene dynamics. For each maturation gene, we computed the time to 10% FC (gene activation/inactivation), time to 50% FC (midpoint of gene activity), and time to 95% FC (gene plateau).

Based on these parameters, we identified 5 gene clusters (**Figure 1H-J, S2, Supplemental Methods**). Of these clusters, three showed time to 10% FC below pseudotime 5. Thus, we termed these clusters early-activating clusters 1,2, and 3 (EA1, EA2, and EA3, respectively). We termed the remaining two clusters late-activating clusters 1 and 2 (LA1 and LA2, respectively). Our computed gene dynamics allowed for a better understanding of CM maturation-related gene changes. Given the prior evidence suggesting the importance of the perinatal window, we expected that a large number of maturation genes would initiate their changes at around pseudotime 20. By contrast, 78% of maturation genes fell into the early-activating categories, and only LA2 (3.4% of maturation genes) showed a time to 10% FC near the onset of the perinatal window. However, apart from EA1, all of the clusters showed time to 95% FC in the pseudotime range [30, 40]. In summary, maturation-related gene changes as a whole likely initiate early. The embryonic/neonatal period is characterized by completion of the EA1 gene program, while the perinatal period is characterized by completion of the EA2, EA3, and LA1 gene programs, as well as initiation and completion of the LA2 gene program. Notably, these dynamics are qualitatively different than those seen in cellular differentiation, which are often driven by “switch-like” genes (28).

Given the differences in dynamics for each of the gene clusters, we were curious to know whether different clusters had different upstream transcriptional regulators. For each cluster, we performed over-representation analysis of TF targets, and further investigated TFs with high fold enrichments across the clusters (**Figure 1K**). This analysis identified several TFs that have been previously implicated in CM maturation, including Srf (29), Err1 (30), Arnt (10), and Mef2 (31, 32), as well as others with previously undescribed function. Interestingly, while some TFs showed stage-specific enrichment, a large number showed enrichment across multiple clusters, despite different transcriptional dynamics. In particular, inferred regulators of EA1 as well as the LA clusters often showed enrichment in other clusters (typically EA2 and EA3). Our results provide a first description of the timing and regulation of CM maturation *in vivo*.

### scRNA-seq recovers maturation trajectory in PSC-CMs

Our next goal was to establish a complementary reference of PSC-CM maturation. In order to directly compare PSC-CMs to our *in vivo* CM trajectory, we generated four separate, isogenic ESC lines from *α*MHC x mTmG mice. We subsequently differentiated these to PSC-CMs using a protocol adapted from previous studies (33–35), with sequential Wnt modulation through defined small molecules (**Figure 2A**). Our rationale was that by using the same parent mouse line, our comparisons could minimize effects of confounding caused by strain/line differences, and isolate biological differences between the *in vivo* and *in vitro* environments.

**Fig. 2.**
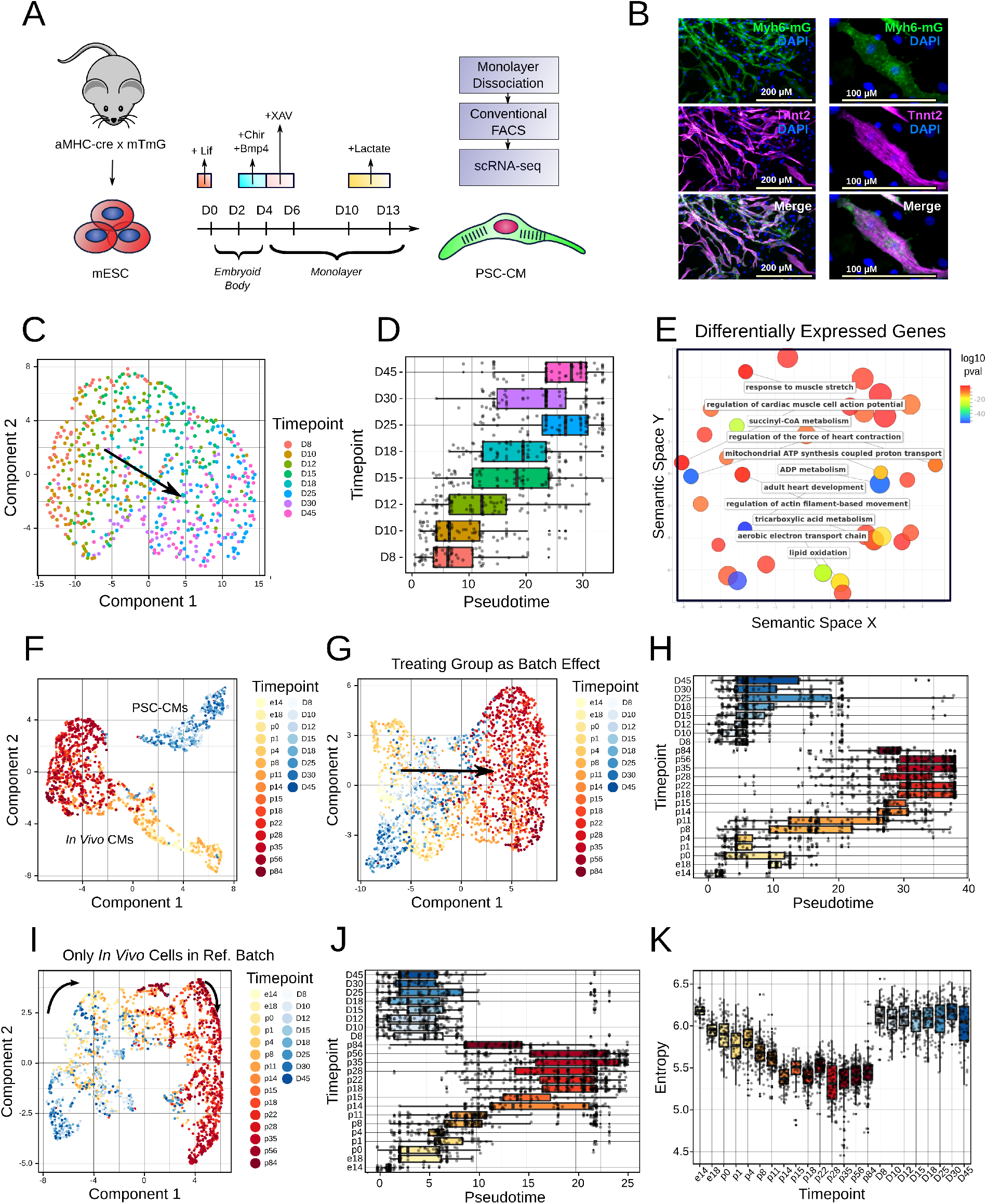
Trajectory reconstruction enables direct comparison of *in vivo* and *in vitro* CM maturation. **A.** Schematic of experimental workflow for generating PSC-CM libraries. **B.** Myh6-mG and Tnnt2 immunofluorescence of PSC-CMs. Right image shows a one sample PSC-CM, with clearly visible sarcomeric structure. **C.** Trajectory inferred by Monocle 3, labeled by timepoint. Arrow indicates the direction of pseudotime. **D.** Pseudotime scores per timepoint for the inferred trajectory. **E.** Gene ontology analysis of genes differentially expressed across the PSC-CM trajectory. **F.** Combined *in vivo* CM and PSC-CM trajectories inferred by Monocle 3, labeled by timepoint. **G.** Combined *in vivo* CM and PSC-CM trajectories with group differences *explicitly* treated as a batch effect by mnnCorrect, inferred by Monocle 3 and labeled by timepoint. **H.** Pseudotime scores per timepoint for the combined trajectory in **2G**. **I.** Combined *in vivo* CM and PSC-CM trajectories with group differences *implicitly* treated as a batch effect by mnnCorrect, inferred by Monocle 3 and labeled by timepoint. **J.** Pseudotime scores per timepoint for the combined trajectory in **2I**. **K.** Transcriptomic entropy for *in vivo* CMs and PSC-CMs across timepoint.

Differentiation of the ESC lines yielded beating GFP^+^ PSC-CMs by Day (D)6.5-7 of differentiation, depending on the line. PSC-CMs additionally displayed other CM markers such as Tnnt2 and displayed sarcomeric structures (**Figure 2B**). We isolated PSC-CMs by conventional FACS from 8 timepoints between D8 and D45 of differentiation for scRNA-seq (**Supplementary Note 1**). Interestingly, at D25, a population of PSC-CMs emerged with light scattering properties that seemed to indicate larger cells (**Figure S3A, Supplemental Methods**). Based on this, we sorted both “large” and “normal” size populations at D25, D30, and D45. We sequenced 660 PSC-CMs to a median depth of ~ 11,400 UMIs per cell and ~3000 genes per cell. SingleCellNet further validated the cells as having CM identity (**Figure S1A**).

As with endogenous CMs, Monocle 3 recovered a unidirectional, non-branching trajectory for PSC-CMs (**Figure 2C, Supplementary Note 3**). Pseudotime progressively but heterogeneously increased over biological time, with a general transition occurring between D10 and D25 of differentiation (**Figure 2D**). These general dynamics were further validated by RNA velocity analysis (**Figure S3B-C**). We observed some, albeit minimal, line-to-line differences in pseudotime progression (**Figure S3D**). Interestingly, there appeared to be no major pseudotime differences between the identified “large” and “normal” cells (**Figure S3E**). This may potentially indicate phenotypic differences in scatter properties that are mediated by non-transcriptional mechanisms. Nevertheless, given these results, we treated these cells as identical for our downstream analyses.

We identified 449 differentially expressed genes over the PSC-CM trajectory. These genes were enriched for terms related to cardiac structure, contractile function, electrophysiology, and metabolism (**Figure 2E**). Taken together, these results validate the successful reconstruction of a PSC-CM maturation trajectory.

### PSC-CMs are transcriptional nearest neighbors to embryonic/neonatal CMs

Having reconstructed trajectories for both *in vivo* and *in vitro* CM maturation, we next sought to align the trajectories to determine how PSC-CMs compared to their *in vivo* counterparts. We combined the two sets of cells and performed trajectory reconstruction (**Supplementary Note 4**). To our surprise, the *in vivo* CM and PSC-CM trajectories remained completely separate, and failed to align (**Figure 2F**). This initial result suggested significant global gene expression differences between the two groups of CMs, despite common genetic background.

To overcome this issue, we made use of properties of mnnCorrect (36), which is the default batch correction algorithm implemented in Monocle 3. mnnCorrect projects cells onto a reference batch by identifying mutual nearest neighbors between the batches, and correcting differences between mutual nearest neighbor pairs. We used mnnCorrect to align the *in vivo* and *in vitro* trajectories in two separate but similar approaches. In the first approach, we set the group (e.g. *in vivo/in vitro*) to be a batch effect to be corrected (**Figure 2G-H**); thus, mnnCorrect *explicitly* looked for mutual nearest neighbors between the groups of cells. In the second approach, we set the reference batch to be a batch containing only *in vivo* CMs (**Figure 2I-J**). This *implicitly* treats the group as a batch effect by again forcing PSC-CMs to be projected onto *in vivo* CMs. Both approaches make the assumption that group differences are either driven by technical artifacts or by biological phenomena that are not of interest. This assumption is likely to be incorrect - indeed, global expression differences may be directly biologically relevant to the poor maturation status of PSC-CMs (which we consider below). Nevertheless, this assumption was appropriate for the initial goal of identifying an approximate alignment between the endogenous CM and PSC-CM trajectories.

Both approaches yielded successful integration of the *in vitro* and *in vivo* CM trajectories (**Figure 2G-J**). Notably, in both approaches, the vast majority of PSC-CMs showed pseudotime score similar to endogenous CMs from e18-p4 (e.g. the embryonic/neonatal period). While some PSC-CMs appeared to enter into the perinatal phase of maturation, nearly no PSC-CMs successfully achieved an adult phenotype. This result is in line with the hypothesis that the previously identified perinatal phase of CM maturation is somehow disrupted in PSC-CMs, leading to their immature phenotype.

As a complementary approach to trajectory reconstruction, we also used our previously developed transcriptomic entropy metric (5) to stage the PSC-CMs. This metric is based on the observation that immature cells display a broader, more promiscuous gene expression profile, which subsequently narrows as the cells mature. Thus, immature CMs will display a high transcriptomic entropy, while mature cells will show a lower transcriptomic entropy. Our previous work optimized this metric to enable direct comparisons across multiple datasets while being robust to technical batch effects. Here, transcriptomic entropy of the PSC-CMs was generally high and corresponded to that of embryonic/neonatal CMs, with very few PSC-CMs demonstrating lower transcriptomic entropy than p8 CMs (**Figure 2K**). These results provide a trajectory reconstruction-independent validation of our above results. Additionally, these results corresponded well with our previous analysis of transcriptomic entropy in human PSC-CMs. Taken together, our data supports the embryonic/neonatal maturation status of PSC-CMs.

### PSC-CMs show global expression differences from endogenous CMs

As a first step in understanding the immature phenotype of PSC-CMs, we sought to determine the global gene expression differences that led to complete separation of the *in vivo* and *in vitro* trajectories. To this end, we tested for differential gene expression between all *in vivo* CMs and PSC-CMs (**Figure 3A**). This identified 2906 differentially expressed genes. 1460/2628 (56%) of our previously identified maturation genes showed global gene expression differences, suggesting significant differences between *in vivo* and *in vitro* CMs. However, one explanation for this could be that this comparison includes mature endogenous CMs, and thus we are simply capturing the immaturity of PSC-CMs. We therefore tested for differential gene expression between early stage CMs (defined as *in vivo* pseudotime ∈ [0, 15]) and PSC-CMs (**Figure 3A**). This yielded 1743 differentially expressed genes, with 1017/2628 (39%) of maturation genes falling into this category. Thus, PSC-CMs showed global expression differences even when compared to endogenous CMs identified as their transcriptional nearest neighbors.

**Fig. 3.**
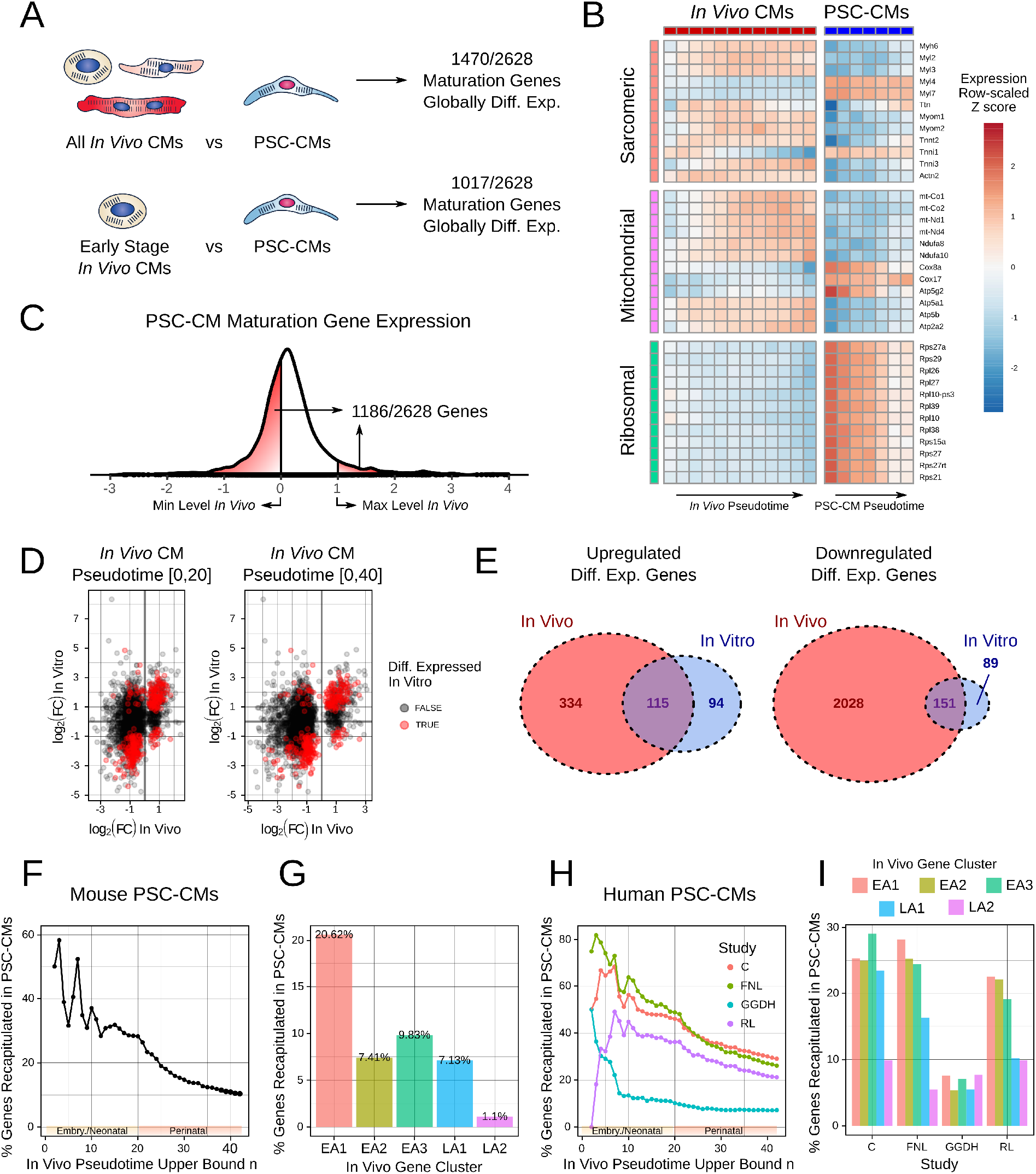
PSC-CM maturation shows both absolute and relative gene expression differences compared to endogenous maturation. **A.** Number of *in vivo* maturation genes that show global expression differences when comparing all *in vivo* CMs to PSC-CMs (top) or early stage CMs (e.g. in pseudotime interval [0, 15]) to PSC-CMs. **B.** Heatmap of candidate sarcomeric, mitochondrial, and ribosomal genes showing global expression differences between *in vivo* and PSC-CMs. Pseudotime units are in increments of 5, based on the individual trajectories for each group (e.g. **Fig 1B** for *in vivo* CMs and **2C** for PSC-CMs). **C.** Histogram of scaled PSC-CM expression levels of *in vivo* maturation genes. We scaled expression levels by setting 0 as the lowest level across *in vivo* pseudotime and 1 as the highest level across in *in vivo* pseudotime. **D.** FC of maturation genes *in vitro* vs *in vivo*, compared for embryonic/neonatal timepoints (e.g. [0, 20], left) and perinatal timepoints (e.g. [0, 40], right). Points are labeled by whether the gene is also differentially expressed in PSC-CMs. **E.** Venn diagrams of differentially expressed genes across pseudotime *in vivo* and *in vitro*. **F.** Percentage of maturation genes in each pseudotime subset (as in **Fig 1D**) correctly differentially regulated in mouse PSC-CMs. **G.** Percentage of genes in each identified gene cluster correctly differentially regulated in mouse PSC-CMs. **H.** Percentage of maturation genes in each pseudotime subset (as in **Fig 1D**) correctly differentially regulated in human PSC-CMs for four studies. **I.** Percentage of genes in each identified gene cluster correctly differentially regulated in human PSC-CMs for four studies.

Gene ontology analysis of globally differentially regulated genes identified terms corresponding to CM maturation (**Figure S4A**). Genes associated with sarcomeric structure, oxidative phosphorylation, fatty acid metabolism, and calcium handling were generally expressed at higher levels *in vivo*, while genes associated with proliferation/cell cycle, stemness, and transcription/translation were higher in PSC-CMs. We further investigated expression differences in major sarcomeric, mitochondrial, and ribosomal genes (**Figure 3B**). In general, sarcomeric proteins and mitochondrial-related transcripts (e.g. mitochondrially-encoded proteins, electron transport proteins, and ATPases) were more highly expressed *in vivo*, while ribosomal protein-coding transcripts were more highly expressed in PSC-CMs. Notably, while PSC-CMs expressed both ventricular myosin light chain genes (*Myl2, Myl3*) and atrial genes (*Myl4, Myl7*), they expressed the former pair at a much lower level than *in vivo* CMs and the latter pair at a much higher level. This is complicated by the fact that endogenous ventricular CMs express *Myl4* and *Myl7* during embryonic stage. Thus, these results may indicate a more “atrial-like” phenotype for PSC-CMs, but they may also reflect a general inadequacy in using these markers to classify CMs (12). Additionally, PSC-CMs showed comparatively higher expression of the immature Troponin I isoform *Tnni1* and lower expression of the mature isoform *Tnni3*.

On the surface, these results seem to reiterate the immature phenotype of PSC-CMs. However, we observed that PSC-CMs do not just demonstrate immature-like gene expression levels. Rather, their absolute gene levels often fall *entirely outside* the spectrum of endogenous development. To quantify this, we scaled all of the maturation genes such that 0 corresponded to the lowest level over *in vivo* maturation, while 1 corresponded to the highest level. We then normalized the average PSC-CM gene expression of the maturation genes by this same scaling. Notably, 1186/2628 genes (45%) fell outside the range of [0, 1] (**Figure 3C**). Strangely, many genes showed lower overall expression levels in PSC-CMs, even if those genes were *downregulated* during endogenous maturation. Gene expression levels for *in vivo* and *in vitro* CMs may thus fall into two entirely different biological scales. These results highlight the challenge of comparing between the *in vivo* and *in vitro* contexts - direct comparison of absolute gene levels may be difficult or even impossible.

### PSC-CMs poorly recapitulate perinatal CM maturation-related gene changes

As expression levels may be on differently calibrated biological scales for *in vivo* and in vitro CMs, direct comparison of expression levels may not yield insights into the dysregulation of maturation *in vitro*. Thus, we proceeded with a different approach by drawing insights from a previous study from our group (37). There, we sought to compare first and second heart field progenitor cells isolated from *in vivo* embryos and *in vitro* precardiac organoids by RNA-seq. While absolute expression levels corresponded poorly between the *in vivo* and *in vitro* samples, the directionality of differences in marker genes between first and cell heart field cells was well preserved. Applying that lesson here, rather than viewing the absolute gene expression levels of the adult CM as the gold standard, we focused instead on the directionality of gene changes from immature to mature CM. We could then see how well these relative changes were recapitulated in PSC-CMs. This approach enabled comparison between the *in vivo* and *in vitro* CM maturation trajectories without the need to directly compare expression levels.

We first computed the FCs for all of the maturation genes across the *in vivo* trajectory, both in the pseudotime interval [0, 20] (encompassing the embryonic/neonatal period) and in the interval [0, 40] (encompassing both the embryonic/neonatal and perinatal periods). We then compared these against the FCs of the maturation genes across the *in vitro* trajectory. Notably, for genes that were differentially expressed both *in vivo* and *in vitro*, FCs were generally comparable (**Figure 3D**). However, only 10% of maturation genes were differentially expressed in the same direction in PSC-CMs (**Figure 3E**). This data supports the failure of PSC-CMs to successfully recapitulate the majority of maturation-related gene changes.

We next used our approach from **Figure 1D** to determine which maturation genes are best recapitulated *in vitro*. We quantified what percent of differentially expressed genes within each pseudotime subset are also differentially expressed in the same direction *in vitro*. While recapitulation in generally poor, genes that become differentially expressed during the embryonic/neonatal period were more likely to be correctly differentially expressed *in vitro* (**Figure 3F**). For example, 37% of genes differentially expressed from [0, 10] *in vivo* are correctly differentially expressed in vitro, and 28% of genes differentially expressed from [0, 20] *in vivo* are captured *in vitro*. However, this number drops to 11% for genes differentially expressed from [0, 40]. The sharp drop-off after *in vivo* pseudotime 20 (corresponding to the start of the perinatal phase) demonstrates that PSC-CM maturation is particularly poor at recapitulating perinatal maturation-related changes.

We further quantified what percentage of genes in the identified *in vivo* clusters are correctly recapitulated *in vitro*. By far, genes from EA1 are best recapitulated in PSC-CMs, while less than 10% of genes in EA2, EA3, and LA1 are correctly differentially expressed *in vitro* (**Figure 3G**). Most strikingly, only 1.1% of LA2 genes are captured *in vitro*. In summary, PSC-CMs undergo a subset of embryonic/neonatal maturation changes, but fail to undergo the perinatal program. This may in turn point to their phenotypic arrest.

### Human PSC-CMs show poor recapitulation of perinatal CM maturation-related gene changes

Given the importance of PSC-CM technology for applications in human health, we next investigated maturation-related changes in human PSC-CMs. In our previous study, we found that, similar to mouse PSC-CMs, human PSC-CMs appear to have a maturation state similar to fetal CMs and appear arrested at the onset of perinatal maturation (5). Here, we focused on four published datasets of CMs generated from induced PSCs - Friedman, Nguyen, and Lukowski et al. (38) (FNL); Churko et al. (39) (C); Gerbin, Grancharova, Donovan-Maye, and Hendershott et al. (40) (GGDH); and Ruan and Liao et al. (41) (RL). We initially aimed to perform trajectory reconstruction with Monocle 3 as done with the mouse PSC-CMs. However, none of the datasets formed a smooth, continuous trajectory but rather showed discrete separation of timepoints (**Figure S4B**). In each of these datasets, samples from individual timepoints were typically prepared as separate batches, and thus batch and timepoint were confounded. Thus, it was difficult to resolve potential batch effects to create an appropriate trajectory. As a work-around, we applied our transcriptomic entropy approach, which we previously showed can also function as a surrogate pseudotime (5). Transcriptomic entropy largely recapitulates similar differentially expressed genes as commonly used trajectory inference methods while being generally resistant to batch effects. Through this method, we identified differentially expressed genes for each of the four datasets.

As with the mouse PSC-CMs, we compared the directionality of changes in maturation genes for the human PSC-CMs against our *in vivo* data. This approach fundamentally assumes that maturation-related changes are comparable across human and mouse. Whether this universally holds requires further assessment; however, in the absence of comprehensive perinatal human CM scRNA-seq data, our assumption served as a useful first approximation. We found that, while several of the human datasets performed better than the mouse PSC-CMs in recapitulation of maturation genes, all showed a sharp drop-off in correct recapitulation at the onset of the perinatal period (**Figure 3H**). Likewise, while the datasets performed better in terms of recapitulating genes in the EA2 and EA3 clusters, the LA1 and LA2 clusters were still relatively poorly captured (**Figure 3I**). The one exception to this observation was the GGDH dataset, which appeared to perform poorly in general, though this may also be for technical reasons owing to the lower depth/sensitivity of that study. As a whole, however, the data indicates that human PSC-CMs similarly fail to undergo perinatal programs associated with CM maturation.

### A network of nine TFs underlies dysregulation of PSC-CM maturation

Given that the poor PSC-CM maturation phenotype can be seen across multiple lines and protocols from several species, we hypothesized that there is a conserved mechanism underlying maturation failure in PSC-CMs. Commonly dysregulated genes across multiple studies may point to a source for disruption of perinatal maturation programs. We thus identified dysregulated genes for the mouse PSC-CMs generated in this study as well as the four literature-obtained human PSC-CM datasets (**Figure 4A**). We used the following criteria to classify a gene as dysregulated: either upregulated *in vivo* but not *in vitro*, or downregulated *in vivo* but not *in vitro* AND expressed *in vitro.* The latter criterion allowed us to eliminate genes downregulated *in vivo* but already not expressed *in vitro*, as these genes are less likely to be relevant to maturation failure. We generated a consensus human list by including genes dysregulated in at least three of the four literature datasets. This list was intersected with the dysregulated genes from the mouse dataset to generate a final list of 550 genes dysregulated across the studies and across species.

**Fig. 4.**
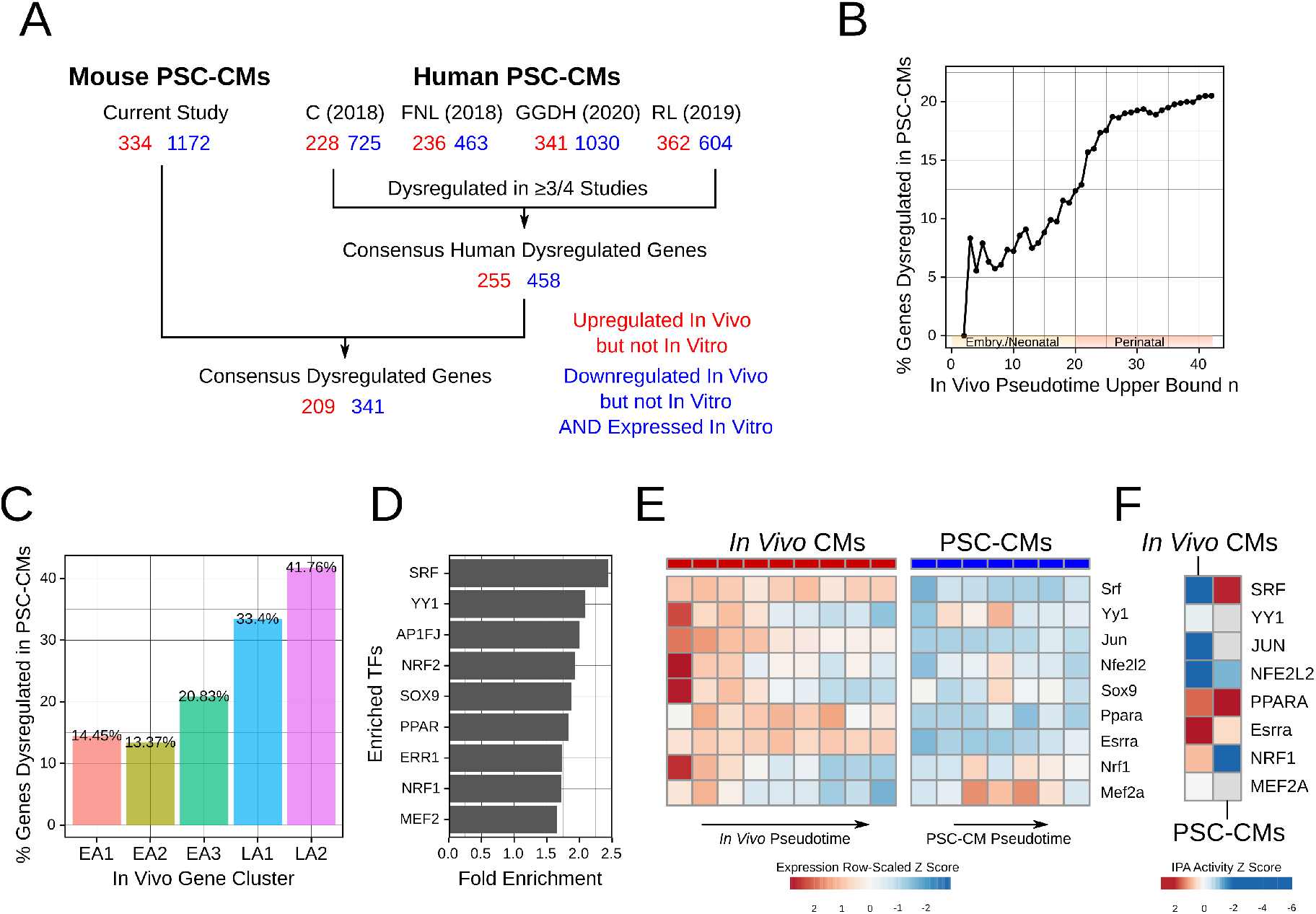
Perinatal maturation programs are dysregulated in PSC-CMs. **A.** Workflow for identifying dysregulated genes in each PSC-CM dataset. **B.** Percentage of maturation genes in each pseudotime subset (as in **Fig 1D**) dysregulated in PSC-CMs. **C.** Percentage of genes in each identified gene cluster dysregulated in PSC-CMs. **D.** TFs whose downstream targets are enriched in the consensus dysregulated gene list. **E.** Scaled expression of identified dysregulated TFs *in vivo* and in PSC-CMs. Pseudotime units are in increments of 5, based on the individual trajectories for each group (e.g. **Fig 1B** for *in vivo* CMs and **2C** for PSC-CMs). The corresponding protein-gene names are: AP1FJ (Jun), NRF2 (Nfe2l2), ERR1 (Esrra). **F.** IPA activity scores for identified dysregulated TFs *in vivo* and in PSC-CMs. Note that SOX9 was not included as an activity score was not assigned by IPA.

The consensus dysregulated gene list was notably enriched for genes differentially expressed during the perinatal period *in vivo* (**Figure 4B**). Similarly, the time to 10% FC, time to 50% FC, and time to 95% were all higher for the dysregulated genes compared to correctly recapitulated genes (**Figure S4C**). Lastly, the consensus dyresgulated gene list was enriched for genes from the LA clusters, with 33% and 42% of genes from LA1 and LA2 dysregulated respectively. These results are in line with our hypothesis that the perinatal period of maturation is particularly disrupted in PSC-CMs. We additionally investigated the chromatin accessibility of the dysregulated genes compared to correctly recapitulated genes by analysis of three previously published ATAC-seq datasets of human PSC-CMs at D15, D25, and D30 (42-44). The percentage of dysregulated genes with peaks in the promoter-TSS region was marginally lower than for correctly recapitulated genes (67% vs 72%) (**Figure S4D**); however, this difference is unlikely to be biologically relevant. This suggests that dysregulated genes show a similar level of chromatin accessibility to correctly recapitulated genes.

Given the similarity in chromatin accessibility, we next aimed to identify upstream factors that could be responsible for gene dysregulation in PSC-CMs. As before, we performed over-representation analysis to identify TFs whose targets are particularly highly represented in the consensus dysregulated gene list. Using affinity propagation to eliminate redundancy (**Supplemental Methods**), we narrowed down the identified TFs to a list of 9 candidate TFs (**Figure 4C**). We refer to these TFs from here as dyregulated maturation TFs. Many of the dysregulated maturation TFs were identified as being important regulators of *in vivo* CM maturation in our above analysis (**Figure 1K**). Additionally, almost all have been previously directly implicated in either CM differentiation, maturation, or disease response (11, 29–32, 45–51). The STRING protein database identified significant connectivity between these TFs (**Figure S4E**), with a protein-protein interaction enrichment p-value of 1.75 × 10^-8^. Thus, the identified TFs likely work as a regulatory network to mediate CM maturation; disruption of this network may underlie maturation failure *in vitro*.

The disruption of our identified TF network could occur at multiple stages, including at the gene expression, protein level, or protein activity levels. As a first step, we investigated the transcriptional levels of each of the dysregulated maturation TFs in our paired *in vivo* and *in vitro* CMs (**Figure 4E**). *In vivo*, all were expressed early in maturation. Subsequently, some decreased in level over the maturation process (*Yy1, Jun, Nfe2l2, Sox9, Nrf1, Mef2a*), while others remained expressed at a relatively constant level (*Srf, Ppara, Essra*). However, in PSC-CMs, nearly all showed much lower levels compared to *in vivo* CMs, particularly at the start of *in vitro* maturation. This supports the possibility of network failure at the gene expression stage. However, as discussed earlier, it is also possible that *in vivo* and *in vitro* CMs have different scales of expression, complicating direct comparison of expression levels. As an alternate comparison, we used Ingenuity Pathway Analysis (IPA) to infer TF activity. IPA infers activity based on fold changes of downstream genes compared against known literature interactions (52). Using IPA, we found that all dysregulated maturation TFs, with the exception of PPARA, showed either weaker or reversed activity over PSC-CM maturation as compared to *in vivo* maturation (**Figure 4F**). Our results support the hypothesis that these dysregulated maturation TFs play a role in the failure of PSC-CMs to undergo perinatal maturation programs.

### Ex vivo perturbations only partially activated dysregulated maturation TF network

A number of cellular and tissue engineering methods have been proposed to improve PSC-CM maturation. However, in the absence of knowledge about *in vivo* maturation or the nature of PSC-CM maturation dysregulation, it has been difficult to assess how biomimetic these methods are. In particular, while these methods may impact some functional characteristics associated with maturing CMs, the mechanism of these changes may be different than endogenous maturation. Our findings here provide a useful for framework for investigating the effects of *ex vivo* perturbations. If a particular perturbation indeed improves PSC-CM maturation in a biomimetic manner, then the direction of gene changes between perturbed vs control tissue should match the direction of changes during *in vivo* maturation.

To this end, we identified six publicly available RNA-seq datasets spanning eight different perturbations of PSC-CMs (53–58) (**Figure 5A**). For each, we identified differentially expressed genes between the provided experimental and control groups, and intersected these genes against our identified maturation genes (**Figure 5B**). Notably, for most of the perturbations, a majority of the identified differentially expressed genes matched the direction of *in vivo* CM maturation. However, a large number (average of ~38% of genes across the 8 studies) were differentially expressed in the opposite direction. While these may represent genes further dysregulated by perturbations, they may also reflect differences in species as well as differences in technical conditions across all of the different studies (e.g. timepoints, isolation protocols, degree of cellular purity). Thus, we focused our analysis on genes that were differentially regulated in the same direction between perturbations and *in vivo* maturation. Of these genes, we found that ~30-50% were already correctly differentially expressed over control PSC-CM maturation in our previously identified datasets (**Figure 5C**). However, ~10-25% of differentially expressed genes came from our consensus dysregulated gene list. Thus, while perturbation methods predominantly impact genes that are already being correctly differentially expressed *in vitro*, they may also “correct” previously dysregulated genes.

**Fig. 5.**
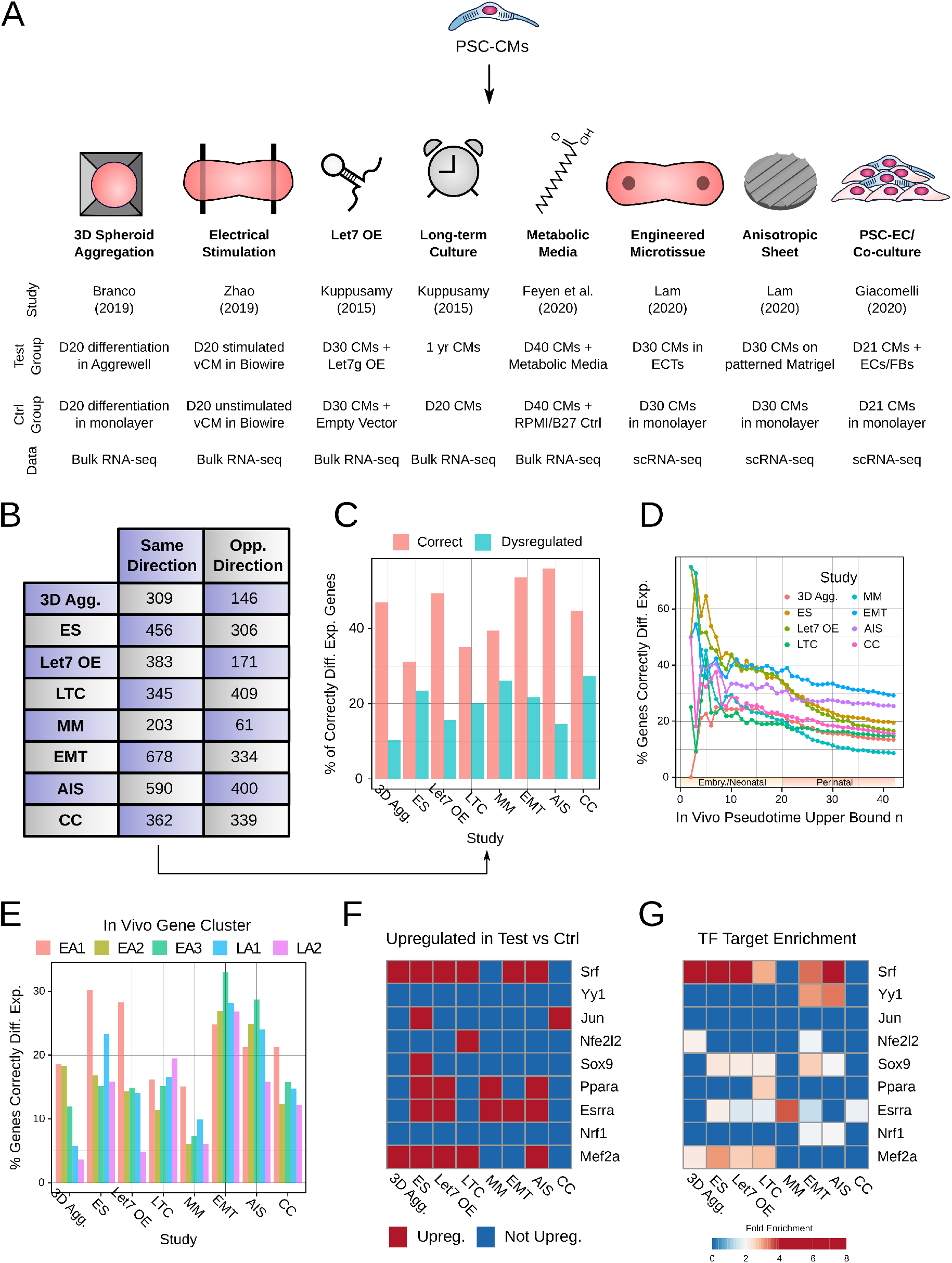
Perturbation methods to engineer PSC-CM maturation activate some, but not all, of the dysregulated maturation TFs. **A.** Characteristics of studies and perturbations analyzed. **B.** Number of maturation genes showing differential change in the same direction following perturbation as *in vivo* maturation. Only genes differentially regulated in the same direction were analyzed in further steps. **C.** Percentage of genes differentially regulated in each study in same direction as *in vivo* CMs that fell into either the “correctly differentially regulated” or “dysregulated” categories in unperturbed PSC-CMs. **D.** Percentage of maturation genes in each pseudotime subset (as in **Fig 1D**) correctly differentially regulated for each perturbation method. **E.** Percentage of genes in each identified gene cluster correctly differentially regulated for each perturbation method. **F.** Plot of which dysregulated maturation TFs are upregulated by each perturbation method. **G.** Heatmap of dysregulated maturation TFs whose downstream targets are enriched in the correctly differentially expressed gene lists for each perturbation method.

For most perturbations, the identified differentially regulated genes were typically enriched in the embryonic/neonatal timepoints (**Figure 5D**), and enriched in genes from EA1 (**Figure 5E**). This suggests that most perturbations predominantly continue to activate early stage gene changes without affecting the dysregulated perinatal phase. However, some perturbations proved to be exceptions to this general rule. In particular, long term culture (LTC) and engineered microtissues (EMT) appeared to show prominent enrichment of LA1 and LA2 compared to EA1 within the same study, potentially indicating that these methods may be better at improving maturation in PSC-CMs.

To further investigate how perturbations affect PSC-CM dysregulation, we looked at changes to our identified dyregulated maturation TFs. We looked at two parameters – whether the TF itself was upregulated by the perturbation (**Figure 5F**), as well as whether downstream targets of the TF were enriched in the differentially expressed genes of that perturbation (**Figure 5G**). The two methods show decent but not complete overlap in results. This may indicate that some methods work through post-transcriptional methods of TF activation, though they may also reflect differences in underlying computational assumptions between the methods. Notably, electrical stimulation (ES) showed the most prominent transcription-level changes to dysregulated maturation TFs, while LTC and EMT showed the largest effect on downstream targets. However, while each method was able to activate some of the dysregulated TFs, no method could completely activate them all. This suggests an inherent limitation in the ability of current *ex vivo* approaches to fully overcome the PSC-CM maturation deficit.

## Discussion

Here, we used scRNA-seq to identify developmental processes associated with CM maturation. In particular, for the first time, we reconstructed a high-quality trajectory of *in vivo* CM maturation with significant sampling of perinatal timepoints. While gene trends associated with CM maturation initiate early (~e18.5), CMs undergo a perinatal phase (p8-p15) during which the rate of transcriptional changes is at its highest and most genes progress to their mature levels. Through trajectory comparison, we found that this perinatal phase is largely not recapitulated in PSC-CMs. We identified a network of nine TFs upstream of dysregulated genes in PSC-CMs; these TFs consistently showed lower expression and disrupted activity *in vitro*. Our study thus provides a transcriptional underpinning for the poor PSC-CM maturation state.

Additionally, by surveying published datasets of *ex vivo* perturbations, we found that no method can fully activate all of the identified TFs. To date, the mechanisms by which *ex vivo* perturbations affect CM biology is often unclear, and maturation is often assessed based on *ad hoc* measurements. Thus, it is possible that these methods work through non-developmentally mimetic mechanisms. We believe that for optimal engineering of PSC-CMs, *in vivo* development must be viewed as the gold standard. Our results emphasize the importance of comparing perturbations directly to endogenous CMs to establish claims of cellular maturation.

CM maturation continues to remain a somewhat poorly understood phenomenon. For example, it is not fully clear what triggers initiate maturation. Our trajectory reconstruction indicates that maturation processes begin *in utero*, in line with previous findings (4, 59). Thus, it is unlikely that birth, and its accompanying hemodynamic and metabolic changes, is the initial driver of CM maturation, though these changes may play a subsequent role. The role of various neuroendocrine cues must be further studied here, as glucocorticoids, thyroid hormone, IGF1, and NRG1 all show spikes in the late embryonic period (60-63). Likewise, while we identified TFs that regulate various clusters of genes through embryonic and perinatal maturation, it is not clear what stimulates activation of these TFs. The analysis of *ex vivo* perturbations, however, can provide biological insight here. For example, perturbations like stretch, electrical stimulation, anisotropic patterning, and others activate maturation-related TFs in partially but not completely overlapping fashion. Thus, maturation TFs may both regulate and be regulated by biophysical changes occurring in the pre- and post-natal heart. Another point of interest is the overlap in upstream TFs across gene clusters with different temporal dynamics. Future studies should more thoroughly identify how common TFs can mediate different expression dynamics, for example through differences in chromatin accessibility, binding affinity, or co-localized TFs.

We identified nine TFs that are likely to underlie maturation failure in PSC-CMs. Interestingly, the dysregulated targets of these TFs show similar chromatin accessibility compared to correctly differentially expressed genes. However, the expression levels of each of the maturation TFs is much lower *in vitro* than *in vivo*. Thus, correcting the expression levels of these TFs provides a putative target for future genetic engineering efforts to improve PSC-CM maturation. Intriguingly, these TFs display significant known interactions with one another, potentially suggestive of a regulatory network for CM maturation. One consequence of this observation is that it is unlikely that upregulation of any one TF alone will induce maturation. Indeed, a recent study found that upregulation of SRF in neonatal CMs led to disruption of maturation (29). Thus, balanced and controlled gene dosages are likely necessary for these TFs to effect maturation. We anticipate that a reprogramming-type approach, featuring concomitant upregulation of multiple dysregulated maturation TFs, will enable future generation of mature PSC-CMs.

Our findings here are based on several assumptions that must be considered critically. In comparing the *in vivo* and *in vitro* trajectories, a fundamental assumption is that all gene changes that occur over endogenous maturation must be associated with or required for completion of maturation. However, many identified differentially expressed genes may be inessential. Indeed, we were surprised to note that many downregulated genes *in vivo* already present with a lower expression *in vitro*. Though we filtered many of these genes from our identified consensus dysregulated list, other genes may similarly be dispensable for maturation. While our study is not equipped to identify such genes, the emergence of improved screening methodologies may help better refine the core gene dynamics of CM maturation in the future (64).

A second assumption was that gene trends across maturation should be generally comparable across multiple datasets. This assumption is reasonable for comparing mouse endogenous CMs and PSC-CMs generated within the same study, as done here. However, whether this assumption holds for our various meta-analyses must be considered more carefully. For one, it is not fully clear whether CM maturation trends must be preserved across species. While recent results from Cardoso-Moreira indicate that developmental trends between species are often divergent (65), the results from Uosaki et al. suggest that CM maturation in particular is relatively well-preserved (66). Both sets of studies were done using bulk cardiac data, and scRNA-seq data is currently unavailable for human perinatal timepoints. Thus, while our approach functions as a useful first approximation of studying PSC-CM dysregulation, there is also a need for better resolution human data to further validate our findings. Likewise, there are generally few well-controlled scRNA-seq studies of *ex vivo* perturbations to improve CM maturation. We anticipate that future datasets can provide better insight into the molecular pathways by which specific perturbations impact CM maturation.

Thirdly, our study primarily focused on transcriptional mechanisms of CM maturation and PSC-CM dysregulation. However, there is plenty of evidence to implicate post-transcriptional processes in CM maturation. For one, we observed that CMs achieve a largely maturation transcriptome by p15-p18. However, protein-level changes continue to occur after this time, and CMs do not achieve their maximal volume until approximately three months of age in mice (67). Moreover, protein-protein interactions at the cell membrane and with the extracellular matrix mediate important changes in postnatal CM biology (68, 69). We expect that the emergence of improved proteomics methods, including single cell proteomics, will enable better resolution of post-transcriptional maturation processes (7, 70). Nevertheless, our results make it clear that CM maturation is associated with large-scale transcriptional changes, and that these changes are not fully recapitulated *in vitro*. Thus, it is likely that transcriptional mechanisms are necessary, though perhaps not sufficient, for successful CM maturation.

Despite these assumptions, we believe this study takes the first steps towards understanding the nature of PSC-CM maturation failure by direct comparison to endogenous developmental processes. These findings can serve as a launch point for future efforts to improve the clinical applicability of PSC-CMs.

## Materials and Methods

All methods, including wet lab and computational methods, can be found in the Supplementary Information. Raw data for the maturation reference can be found on GEO at GSE164591. Code to generate figures in this manuscript as well as the counts tables for the datasets analyzed in this manuscript can be found on Github at https://github.com/skannan4/cm-dysregulation, and further files can be found on Synapse (https://www.synapse.org/#!Synapse:syn23667436/files/).

## ACKNOWLEDGMENTS

We thank Dr. Deborah Andrew for allowing us to use her group’s COPAS LP-FACS instrument. Conventional FACS experiments were done through the Bloomberg Flow Cytometry and Immunology Core/Cell Sorting Facility with the help of Dr. Hao Zhang. Sequencing experiments were done through Novogene with the help of Sanya Pal, Payton Hovey, Dr. Mary Grantham, and Victor Popoca. Additional assistance was provided by the Johns Hopkins Transcriptomics and Deep Sequencing Core, with the help of Dr. Haiping Hao, Linda Orzolek, Dr. Jasmeet Sethi, and Kelly Laughlin. We additionally thank Dr. Patrick Cahan, Yuqi Tan, Emily Su, Taibo Li, and Elaine Zhelan Chen for helpful comments in preparing this manuscript. This work was supported by grants from NICHD/NIH, AHA, and MSCRF.

## Supporting Information Text

The purpose of our supplementary materials section is to provide detailed information about our study that was not included in the main manuscript text. We aspire to standards of data reproducibility and availability. To this end, all of the sequencing data for this study can be found on GEO with accession number GSE164591. Additionally, the code to reproduce all of the figures in the manuscript is available on Github at https://github.com/skannan4/cm-dysregulation. Lastly, we have made an R workspace available on Synapse (https://www.synapse.org/#!Synapse:syn23667436/files/) that contains many of the data tables pre-loaded for our analysis. If there are other materials that could facilitate re-analysis or exploration, we would be glad to provide them on inquiry.

The supplementary materials here contain methods and notes. The supplementary methods provide details for our mouse handling, cell isolation, and sequencing. We also provide rough information on the computational methods used, though we direct readers directly to the code for more involved details. The supplementary notes delve into our library preparation design as well as an overview of how our trajectories were reconstructed in Monocle 3. Again, for more detail, we encourage readers to directly examine the code.

## Supplementary Methods

### Mice and PSC-CM lines

To generate mice for our reference dataset, we crossed B6.FVB-Tg(Myh6-cre)2182Mds/J mice (*α*MHC-cre, Jackson Laboratory, Stock No. 011038) with B6.129(Cg)-Gt(ROSA)26Sor^tm4(ACTB-tdTomato,-EGFP)Luo/J^ (mTmG, Jackson Laboratory, Stock No. 007676). Both mice have C57BL/6J congenic background. To generate mESCs for *in vitro* studies, mice of the above strain were crossed and observed for the presence of a vaginal plug (considered embryonic day 0.5). On embryonic day 3.5, blastocysts were flushed, isolated, and maintained in pre-gelatinized 96 well plates with 2i media. Four genotype-verified mESC lines were expanded and tested for differentiation capability to the cardiomyocyte lineage. All animals were maintained compliant to protocols by the Johns Hopkins Animal Care and Use Committee.

### CM Isolation

For isolation of CMs from e14-p4 timepoints, we used the neonatal cardiomyocyte isolation kit from Miltenyi Biotec in conjunction with the gentleMACS Dissociator. For later timepoints, we performed Langendorff isolation of CMs. We prepared the following buffers:

1. Perfusion buffer: 120 mM NaCl, 5.4 mM KCl, 1.2 mM NaH_2_PO_4_, 20 mM NaHCO_3_, 5.5 mM glucose, 5 mM BDM, 5 mM Taurine, and 1 mM MgCl_2_, adjusted to pH 7.4
2. Digestion buffer: 40 mL Perfusion buffer plus 35.8 mg Collagenase Type II (Worthington CLS-2), 3 mg Protease (Sigma P5147)
3. Tyrode’s buffer: 140 mM NaCl, 5 mM KCl, 10 mM HEPES, 5.5 mM glucose, and 1 mM MgCl_2_, adjusted to pH 7.4

We used a horizontal (i.e. non-hanging) Langendorff apparatus with a chamber filled with perfusion buffer. To perform isolation, we first performed isofluorane anaesthesia on non-heparinized mice. Mice were observed until clearly anaesthetized and unresponsive to toe pinch, and subsequently euthanized by cervical dislocation. The heart was then rapidly excised from the chest and cannulated to the Langendorff apparatus. Flow time and rate of flow were dependent on the age of the mouse and were typically judged based on completeness of digestion to touch. Subsequently, the left ventricular free wall was excised and minced. We filtered isolated cells through a 100 *μ*M screen to eliminate large tissue chunks, spun down at 800 RPM for 1 minute (Eppendorf centrifuge 5702), and resuspended cells in 10 mL Tyrode’s buffer.

### PSC-CM Differentiation

Ûur PSC-CM differentiation protocol was adapted from multiple previously published protocols. The day prior to initiation of differentiation protocol (D-1), cells were changed to expansion media with 1000 U Lif/mL with no CHIR99021 or PD0325901. On Day 0 of differentiation, cells were dissociated with TrypLE (Thermo Fisher) and suspended in a serum-free differentiation media (SFD). SFD was composed of ¾ volume IMDM to ¼ volume Ham’s F12 media, with 0.5% v/v N2 supplement (Gibco), 1% v/v B27 without retinoic acid (Gibco), 0.5% v/v BSA (Invitrogen) in PBS, 0.75% v/v glutamine (Gibco), 0.75% v/v penicillin-streptomysin (Gibco), 50 *μ*g/mL ascorbic acid (Sigma), and 0.039 *μ*l/mL 1-thiogylcerol. For the first four days of differentiation, differentiating cells were maintained as embryoid bodies. On Day 2, media was replaced with fresh SFD plus 3*μ*M CHIR99021 (Selleckchem) and 2.5 ng/mL BMP4 (R&D Systems). On Day 4, cells were dissociated with TrypLE and replated as a confluent monolayer on gelatin-coated flasks. At this time, cells were cultured with SFD with 0.1% v/v XAV939. On Day 6, the media was changed to SFD. From Day 10 to Day 13, lactate selection was performed by culturing the cells in DMEM without glucose plus lactate. Subsequently, the cells were cultured in SFD, with media changes every two days.

### FACS for Single CM and PSC-CM Isolation

For isolating endogenous CMs, we used LP-FACs. We have detailed our LP-FACS approach previously (1). We reproduce our methods here. We utilized a COPAS SELECT instrument (Union Biometrica). The COPAS SELECT was updated and rebranded as the FP-500, but the protocol here study does not use the new features and thus the two are functionally indistinguishable. We optimized sorting for cardiomyocytes by using a sort delay of 8 and sort width of 6. Additionally, we used the following fluorescence settings: ext gain 50, green gain 200, yellow gain 200, red gain 255, extension integral gain 50, green integral gain 200, yellow integral gain 200, red integral gain 255, green PMT 800, yellow PMT 800, red PMT 1100. Coincidence check was selected to ensure proper single event sorting. We typically flowed cells between 20 - 60 events/second. We maintained cells in Tyrode’s buffer during the sort and sorted them into prepared collection plates for mcSCRB-seq library prep. To run the machine, we used ClearSort Sheath Fluid (Sony, Lot 1218L345).

PSC-CMs, unlike endogenous CMs, do not retain their shape when dissociated and instead round up. Thus, they fall below the recommended range of sorting through LP-FACS, but can be readily sorted through conventional FACS. Here, PSC-CMs were dissociated from culture flasks with TrypLE (5-10 minutes, depending on the timepoint). Cells were strained to remove clumps and clusters and subsequently sorted into prepared collection plates on either a MoFlo Legacy or MoFlo XDP. We isolated healthy singlets by first gating on forward and side scatter, followed by forward scatter and pulse width. We used a propidium iodide stain for further isolation of healthy cells, sorting out GFP^+^ /PI^-^ cells. Interestingly, from D25 onwards, we observed that GFP^+^/PI^-^ cells appeared to split into two populations based on PI autofluorescence (despite being PI^-^). When analyzed, the high autofluorescence cells also appeared to have higher side scatter and pulse width (**Figure S3A**). Upon sorting, these cells also appeared to be visually larger under the microscope. Given this, we sorted cells from both the “normal” and “larger” populations at D25, D30, and D45.

### scRNA-seq Library Preparation and Sequencing

We performed sequencing using the mcSCRB-seq protocol (2). The protocol has been described at protocols.io at <u>dx.doi.org/10.17504/protocols.io.p9kdr4w</u>. Information about the library design is provided in the **Supplementary Notes**; more detailed metadata is also available on Synapse. We sequenced the final, pooled library on a NovaSeqS4 as 150-base pair paired-end reads. We subsequently demultiplexed into two files such that the read 1 file contains the 8 base pair i7 tag, 6 base pair cell barcode, and 8 base pair UMI; read 2 contains the 150 base pair cDNA read. We have provided these final demultiplexed reads to GEO at accession GSE164591. However, if the original paired-end reads from the sequencer are desired, we are happy to provide on request. Mapping of the data was done with kallisto|bustools (0.46.2) (3), using an index generated from the CellRanger mouse reference concatenated with the ERCC spike-in sequences. For RNA velocity, mapping was done with kallisto|bustools using special indices with intronic and exonic sequences respectively from GRCm38.98.

### Computational Analysis

Most analyses performed in the paper were done in R (with the exception of RNA velocity, done in Python); code to reproduce the figures can be found at our Github (https://github.com/skannan4/cm-dysregulation). We encourage readers to look directly to the code for specific technical details about our method, and we are of course happy to answer additional questions on request. However, here, we briefly annotate methods used throughout the manuscript.

### General Quality Control. (Figure S1A)

Quality control continues to be a major issue in scRNA-seq analysis. Poor quality cells can confound analyses and need to be removed. In this manuscript, we used the general approach to quality control established in our previous work (4). We used three parameters: percent of reads going to the top 5 genes, depth, and CM identity. For the first two parameters, we normalized the computed metric against the median value for that timepoint, since both metrics will inherently vary as CMs mature. Our thresholds were top5 norm < 1.8 and depth norm > −0.7. These are somewhat more permissive than the thresholds we set in our previous work; however, in that study, we were working with many datasets, including droplet datasets were poor quality cells are more abundant. We found empirically that these relaxed thresholds were sufficient for our higher quality plate-based data. For CM identity, we used the singleCellNet package (0.1.0) (5), compared against the Tabula Muris reference (6). We selected all cells whose highest scoring identity was “cardiac muscle cell.”

### Trajectory Reconstruction. (Figures 1B, 1C, 2C, 2F, 2G, 2H, 2I, 2J, 2K)

For trajectory reconstruction, we used Monocle 3 (0.2.3.3) (7). Monocle 3 uses fastMNN based on the batchelor package (1.2.4), which in turn is based on mnnCorrect (8). We discuss some specifics of trajectory reconstruction (e.g. number of principal components used, batch effect removal) in the **Supplementary Notes**. More specific details such as trajectory start points and graph settings can be found in the code.

### Differential Gene Expression and Gene Ontology Analysis. (Figures 1D, 2E, 3A-I, 5A-E, S1B, S2A-D, S4A-B)

Differential gene expression analysis for single cell datasets was done in Monocle 3. We typically set a cutoff such that testing was only done in genes expressed in at least 25% of cells; we used this as our cutoff for whether a gene was “expressed” in a given cell group. A Benjamini-Hochberg-adjusted p-value threshold of q < 0.05 was used to determine significance.

For bulk datasets, as in **Figure 5**, we instead used DESeq2 (1.26.0) (9), typically setting the design to compare between perturbation and control. We used the Benjamini-Hochberg-adjusted p-value threshold of q < 0.05 to determine significance. However, because bulk samples can detect genes with higher sensitivity, we additionally used a fold change threshold of | log2(FC)| ≥ 0.5.

For Gene Ontology analysis, we predominantly used the resource at the Gene Ontology website (http://geneontology.org/), which in turn links to the PANTHER classification system (10). To more readily visualize Gene Ontology terms while removing redundancy, we selected the top 150 terms by enrichment and input into REVIGO (11). The exception to this workflow was in **Figure S2**, where we instead used WebGestalt 2019 with the following settings: biological processes terms, size limit 850, top 25 terms, weighted cover set expecting 10. Our rationale for using this second method was that the terms provided were more readily condensed for easier broad visualization.

### RNA Velocity. (Figures 1E, 1F, S3B, S3C)

Intronic and exonic matrices were loaded into Python for analysis with scvelo (0.2.2) (12). Rather than use scvelo’s method to identify dispersed genes, we used the identified differentially expressed genes associated with each trajectory. Likewise, we used the computed aligned principal components from Monocle 3 in scvelo. The full dynamical mode was computed for gene velocities, and this was projected onto the Monocle 3 trajectory.

### Gene Clustering. (Figures 1I, IJ)

We detail the approach to compute the time to 10% FC, time to 50% FC, and time to 95% FC in the manuscript. We identified clusters by performing k-means clustering for the genes on these three parameters. Our approach to determining the appropriate number of clusters was empirical - we added clusters so long as clusters with new properties emerged, and stopped once adding a cluster resulted in two clusters with near identical properties. This led to an optimal k = 5.

### TF Analysis. (Figures 1K, 4D, 4F, 5G)

TF enrichment analysis was done using WebGestalt 2019, performing over-representation using the “transcription factor target” database. We started by selecting all TFs with FDR < 0.05. This generally produced a list with significant redundancy; we used different approaches to handle the redundancy depending on the situation. In **Figure 1K**, we first took all TFs identified as enriched for each cluster (or group of clusters, in the case of LA1 + LA2). We manually aggregated redundant TFs by first labeling TFs into supergroups of interest (for example, MEF2 and RSRFC4 could be combined), and then selecting the term with the highest enrichment. We then selected the top 25 for visualization in the figure by selecting the TFs with the top summed enrichments across all clusters - this inherently picked TFs represented across multiple clusters, though we note that nearly no TFs were identified in only EA1 or LA. For **Figure 4D**, we were instead interested in top candidates. Thus, we used the affinity propagation method in WebGestalt, which essentially clusters TFs based on their downstream targets. For each cluster, we selected as representative the TF with the highest expression in *in vivo* CMs (which typically matched the TF selected by WebGestalt). For **Figure 5G**, we selectively used fold enrichments for our TFs of interest, selecting the term with the highest enrichment ratio. Activity analysis was done in IPA (13), using the list of differentially expressed genes and the appropriate fold change for each group.

### ATAC-seq Analysis. (Figure S4D)

Our goal with the ATAC-seq datasets was to quantify the percentage of genes in a list of interest with peaks at the promoter-TSS region. In general, we used the peak calling settings from the original manuscripts, typically with q-value < 0.05 as a threshold. For Liu et al. and Bertero and Fields et al., the appropriate data was downloaded from GEO as BED or narrowPeaks output files from MACS2 (14, 15). We annotated peaks using HOMER (4.11.1) (16), and subsequently filtered peaks with annotation “promoter-TSS” For Greenwald and Li et al., data was available as bigWig files. We therefore converted to WIG (bigWigToWig) followed by conversion to BED (wig2bed, 2.4.38) (17). We then called peaks from the BED file using MACS2 (2.2.7.1) using the peak calling settings from the original manuscript as shown on GEO. We then annotated peaks as above with HOMER. The choice of HOMER genome matched the original study, e.g. hg19 for Liu et al. and Greenwald and Li et al., and hg38 for Bertero and Fields et al.

## Supplementary Notes

### Supplementary Note 1: Library Design

Batch effects in library design can play a critical role in the interpretation of results (18). Indeed, while sophisticated batch correction methods are now available, they are generally at their most efficient for carefully balanced batch designs, and indeed some confounded batch designs may never be well-resolved even with correction algorithms. Based on the parameters of our sequencing approach and limitations in sample isolation and preparation, we aimed to balance our batches as carefully as possible.

In this study, we used the mcSCRB-seq protocol (2) to generate sequencing libraries. Briefly, mcSCRB-seq is a plate-based UMI protocol. Barcoding of cells arises from two steps - the introduction of a cell-specific barcode during reverse transcription, and through the i7 barcode added after tagmentation. The first barcode is added on prior to pooling of cells, while the i7 is added subsequent to pooling. However, the primer used during reverse transcription is expensive, and thus we had access to a 96-barcode set of primers. This inherently limited the number of cells we could pool per initial library to 96 cells. We were able to generate additional multiplexing power through use of numerous i7 primers, which are comparatively cheaper. However, the need for many libraries necessarily introduced potential batch effects. Additionally, once pooled, libraries were processed in batches at different times based on when the samples were isolated as well as limitations in the number of samples we could simultaneously process. We aimed to balance our libraries and batches such that any technical effects could be subsequently removed. We used the following design:

#### Batch 1

Batch 1 was composed of 12 libraries (of 96 cells each), entirely comprised of *in vivo* CMs. The samples were isolated between Dec. 2018 and Feb. 2019, and processed for sequencing in May 2019. Each library is composed of a mix of e14, e18, p0, p4, p8, p11, p14, p18, p22, p28, p35, and p56 CMs, with all timepoints represented in each library. The full preparation for each of the 12 libraries was completed at the same time.

#### Batch 2

Batch 2 was comprised of 10 libraries, entirely comprised of *in vivo* CMs. The samples were isolated between Jun. 2019 and Aug. 2019, and processed for sequencing in Nov. 2019. Each library is composed of a mix of p0, p1, p4, p8, p11, p14 or p15, p18, p22, p28, p35, p56, and p84 CMs, with all timepoints represented in each library. The full preparation for each of the 10 libraries was completed at the same time.

#### Batch 3

Batch 3 was comprised of 10 libraries. The majority of cells were *in vitro* CMs; however, we spiked *in vivo* CMs into each library such that subsequently, we could rule out the possibility that any *in vivo* vs *in vitro* difference was entirely due to batch effects. The samples were isolated between Sept. 2019 and Oct. 2019, and processed for sequencing in Nov. 2019 (albeit independently from Batch 2). Each library is composed of a mix of D8, D10, D12, D15, D18, D25, D30, D45 PSC-CMs and either p56 or p84 CMs, with all timepoints represented in each library.

Thus, based on our design, despite the likelihood of technical effects arising both within and between batch, we anticipated that the significant overlap between libraries and between batches would enable us to better regress out technical differences while preserving biological variation. We pooled all of the individual libraries at equimolar amounts to produce one final sequencing library, which sequenced on one sequencing lane (thereby eliminating lane-to-lane effects as a potential technical confounder).

### Supplementary Note 2: Generating the Endogenous CM Trajectory

We performed reconstruction of the *in vivo* CM trajectory in Monocle 3 using the standard approach, with 5 principal components used in the preprocessing step. We selected this number of principal components somewhat empirically, though we found that results were generally consistent when varying that parameter. Initial reconstruction, with no correction approach, yielded a trajectory that demonstrated timepoint-based differences, but also showed clear separation based on batch (**Figure S5A**). This was somewhat expected, given that our batches were prepared at different times. However, we found that the deficit could be readily corrected using mnnCorrect. We tried setting both “library” and “batch” as the alignment group in mnnCorrect, and found that library-correction yielded slightly better results (likely owing to the simultaneous removal of within-batch differences). This correction yielded the trajectory shown in **Figure 1B**. Comparison of pseudotimes across batches shows generally good concordance (**Figure S5B**). However, we note that some batch differences still remain; indeed, one of our rationales for focusing on pseudotime ∈ [0, 42] was that we suspected that very late stage pseudotime differences (e.g. ∈ [50, 60]) may be more driven by technical differences rather than biological.

### Supplementary Note 3: Generating the PSC-CM Trajectory

We initially used the same approach as above to generate the PSC-CM trajectory, using 3 principal components and “library” as the mnnCorrect alignment group (as there is only one *in vitro* batch). However, the yielded trajectory showed no correspondence with timepoint (**Figure S5C**). Our initial investigation found that while some CM-related genes varied across this trajectory, it was also largely driven by transcriptional noise unrelated to our phenomenon of interest.

We were not entirely surprised by this result as we have previously observed similar occurrences in other datasets, in both Monocle 2 and 3. We generally suspect that a totally unbiased upstream approach (using either dpFeature as in Monocle 2 (19) or UMAP as in Monocle 3 (7)) can sometimes hone in on processes outside of those of biological interest. Therefore, we applied a semi-supervised approach that we have previously used with Monocle 2 (20). We first identified differentially expressed genes between the first (D8) and last (D45) PSC-CM timepoints. We then used only these genes to perform the preprocessing steps (again, with 3 principal components and “library” as an alignment group). This yielded the trajectory in **Figure 2C**, which shows both timepoint-related differences and gene expression changes relevant to maturation. Notably, we identified more genes differentially expressed across this trajectory than we did between D8 and D45 alone. This likely occurred due to both an increased number of cells for comparison and better modeling enabled by trajectory reconstuction.

As an aside, the semi-supervised approach utilized above also works for the *in vivo* trajectory, yielding largely similar results. We didn’t use it for our *in vivo* trajectory simply because there was no need to, as timepoint differences were the primary driver of biological variance in that system.

### Supplementary Note 4: *In vivo* and *in vitro* trajectory alignment

Our first approach to trajectory alignment involved simply combining all of the *in vivo* and *in vitro* cells and performing reconstruction as above. By setting one of the libraries from Batch 3 (e.g. containing both *in vivo* and *in vitro* CMs) as the reference batch, we could use mnnCorrect to correct between-library differences without removing any biological differences between *in vivo* and *in vitro* groups. We used 15 principal components for dimensionality reduction (again, selected empirically, using more PCs to account for more cells in the combined trajectory) and “library” as the alignment group. This produced the trajectory in **Figure 2F**, wherein the *in vivo* and *in vitro* CMs completely separated. We could validate that these differences are largely biological as the *in vivo* CMs in Batch 3 align with other in vivo CMs rather than other cells in Batch 3 (**Figure S5D**).

To force alignment, we used two approaches as discussed in the main manuscript. In the first approach, we first preprocessed with 15 principal components, followed by mnnCorrect alignment across “library.” We then did a second mnnCorrect alignment step across “group.” This enables nonlinear correction of both between-library differences and between-group differences. This yielded **Figure 2G**. In the second approach, we performed the exact same approach as for **Figure 2F**, but instead set a library from Batch 1 (e.g. only *in vivo* cells) as the reference batch. This yielded **Figure 2I**. While both approaches yield comparable approaches, the first seemed to handle true batch effects more effectively (as can be seen by the position of p84 cells from Batch 3).

**Fig. S1.**
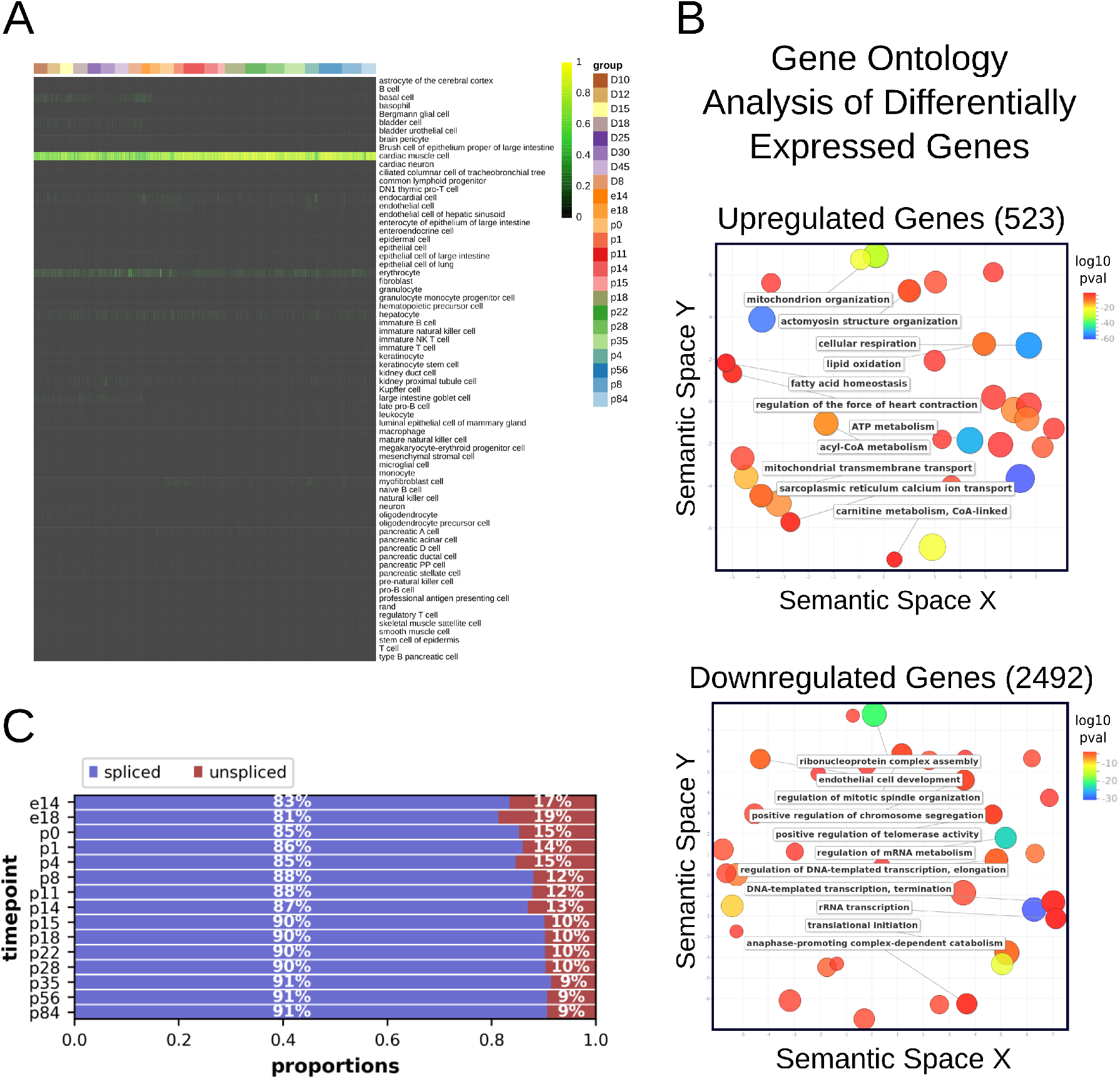
scRNA-seq accurately samples an *in vivo* CM maturation process. **A.** Heatmap of cell classifications of *in vivo* and *in vitro* CMs used in the study by SingleCellNet. The Tabula Muris was used as the background reference for training the classifier. **B.** Gene ontology analysis of genes differentially expressed over the inferred CM maturation trajectory in **Fig. 1B**. **C.** Percentage of counts mapped to spliced and unspliced transcripts for each timepoint.

**Fig. S2.**
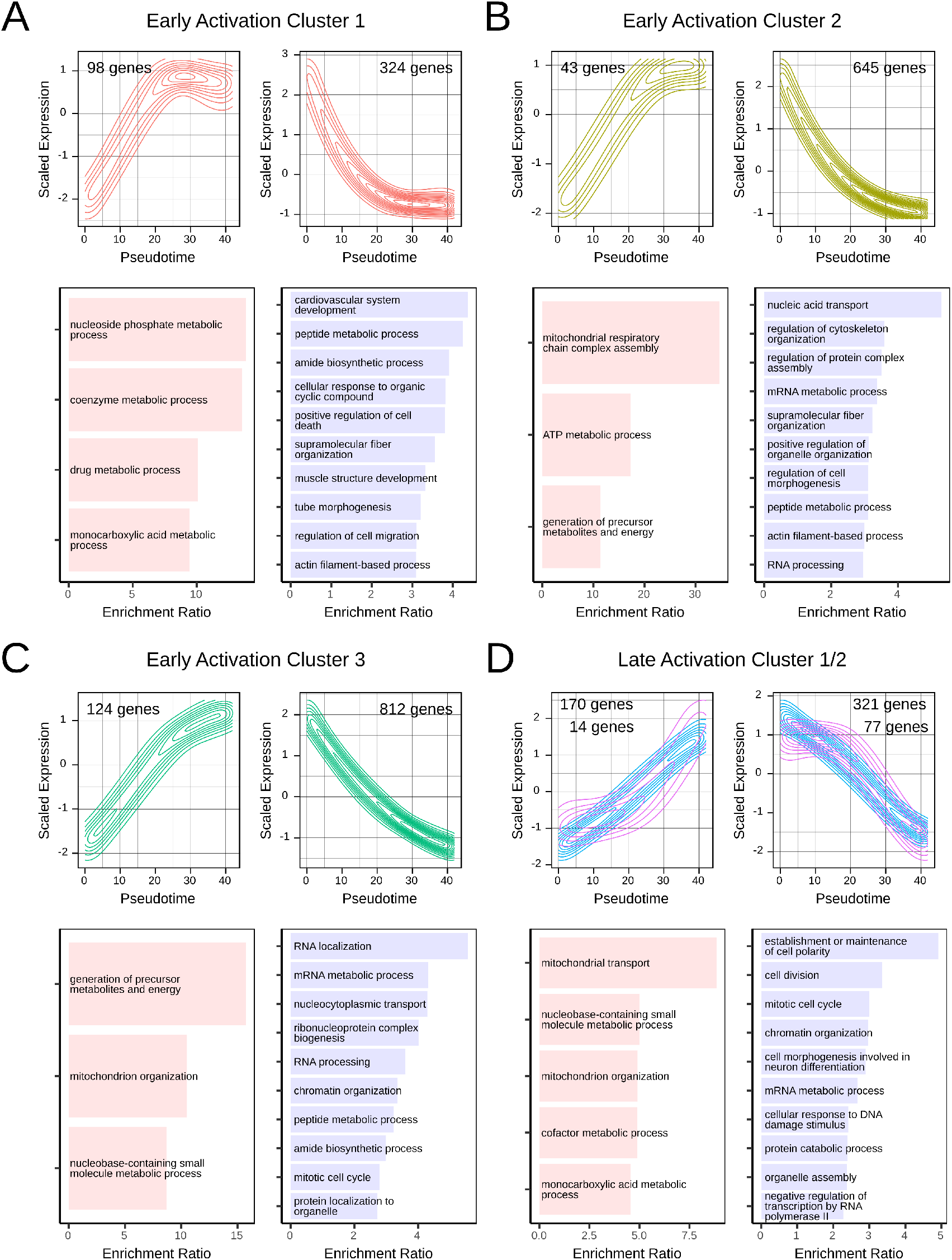
Gene ontology analysis captures biological processes occurring within each temporally-defined gene cluster. **A-D.** Gene ontology analysis for upregulated and downregulated genes in each identified cluster. As cluster LA2 has relatively few genes, we combined with LA1 for analysis.

**Fig. S3.**
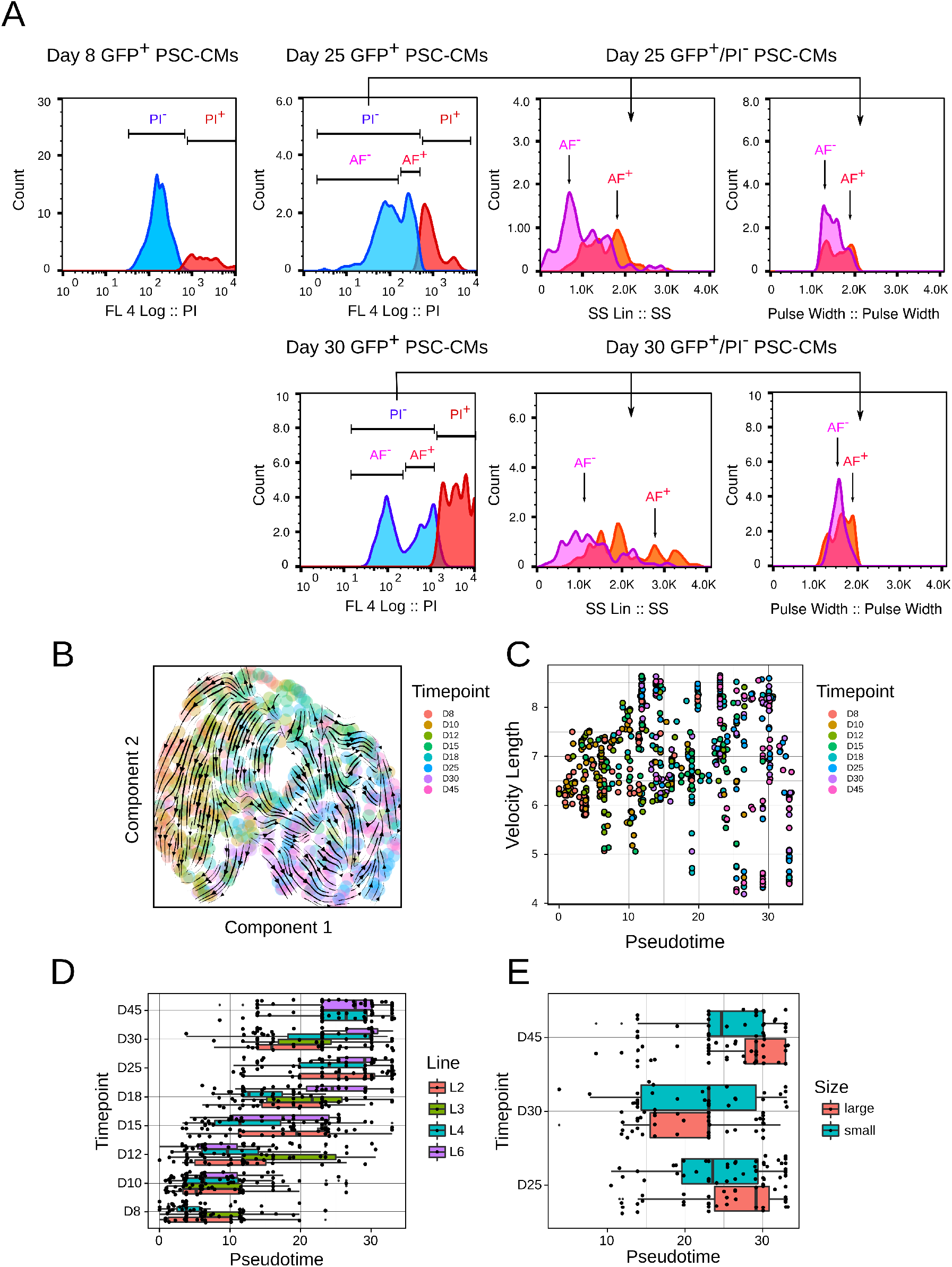
scRNA-seq captures a maturation process in PSC-CMs. **A.** Flow cytometry parameters for GFP^+^ PSC-CMs at D8, D25, and D30 of sorting. Data shown for one cell line (Line 2) but comparable results were seen in other lines. AF = autofluorescence. **B.** RNA velocity stream plot projected onto inferred PSC-CM trajectory, labeled by timepoint. **C.** RNA velocity length across Monocle 3-inferred pseudotimes for PSC-CMs. **D.** Pseudotime scores per timepoint for the inferred trajectory, further labeled by cell line. **E.** Pseudotime scores per timepoint for the inferred trajectory, further labeled by putative size category (as in **S3A**).

**Fig. S4.**
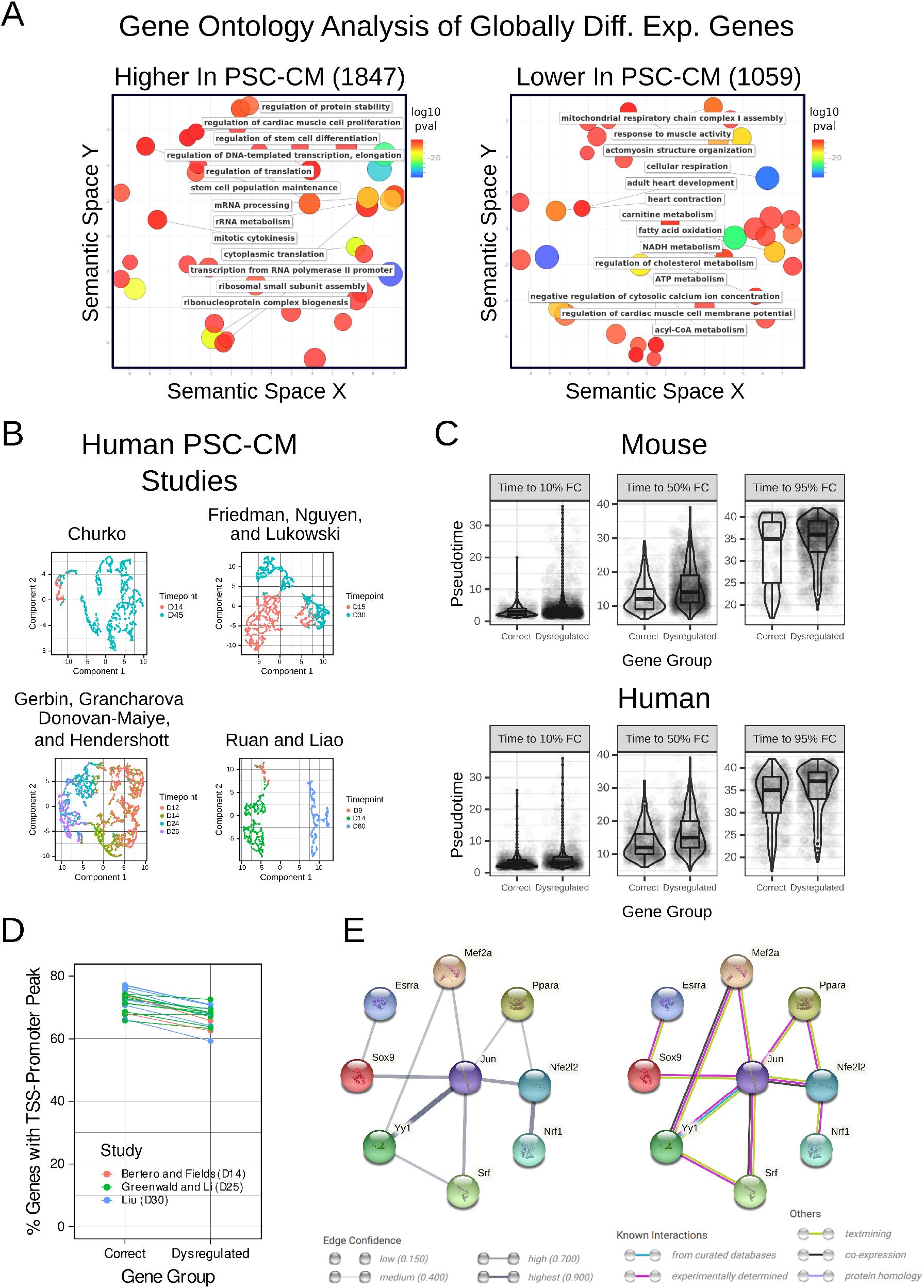
PSC-CMs show dysregulated genes compared to *in vivo* CM maturation. **A.** Gene ontology analysis of all genes globally differentially expressed between endogenous CMs (at all timepoints) and PSC-CMs. **B.** Dimensionality reduction of four human PSC-CM studies from the literature, labeled by timepoint. Complete separation of timepoints indicates a potential batch effect. **C.** Comparison of gene dynamics parameters between correctly regulated and differentially regulated genes in mouse and human PSC-CMs. For human PSC-CMs, correctly regulated genes were defined as genes with correct direction of differential expression in at least two studies. All comparisons were statistically significant by t-test with *α* = 0.05. **D.** Percentage of correctly regulated or dysregulated genes in human PSC-CMs with ATAC-seq peak in the promoter-TSS region. Points are labeled by study, each of which encompasses a different timepoint, and points are connected by sample. **E.** STRING output networks of the identified dysregulated maturation TFs. Left shows connectivity by edge confidence, while right shows connectivity by type of evidence for interaction.

**Fig. S5.**
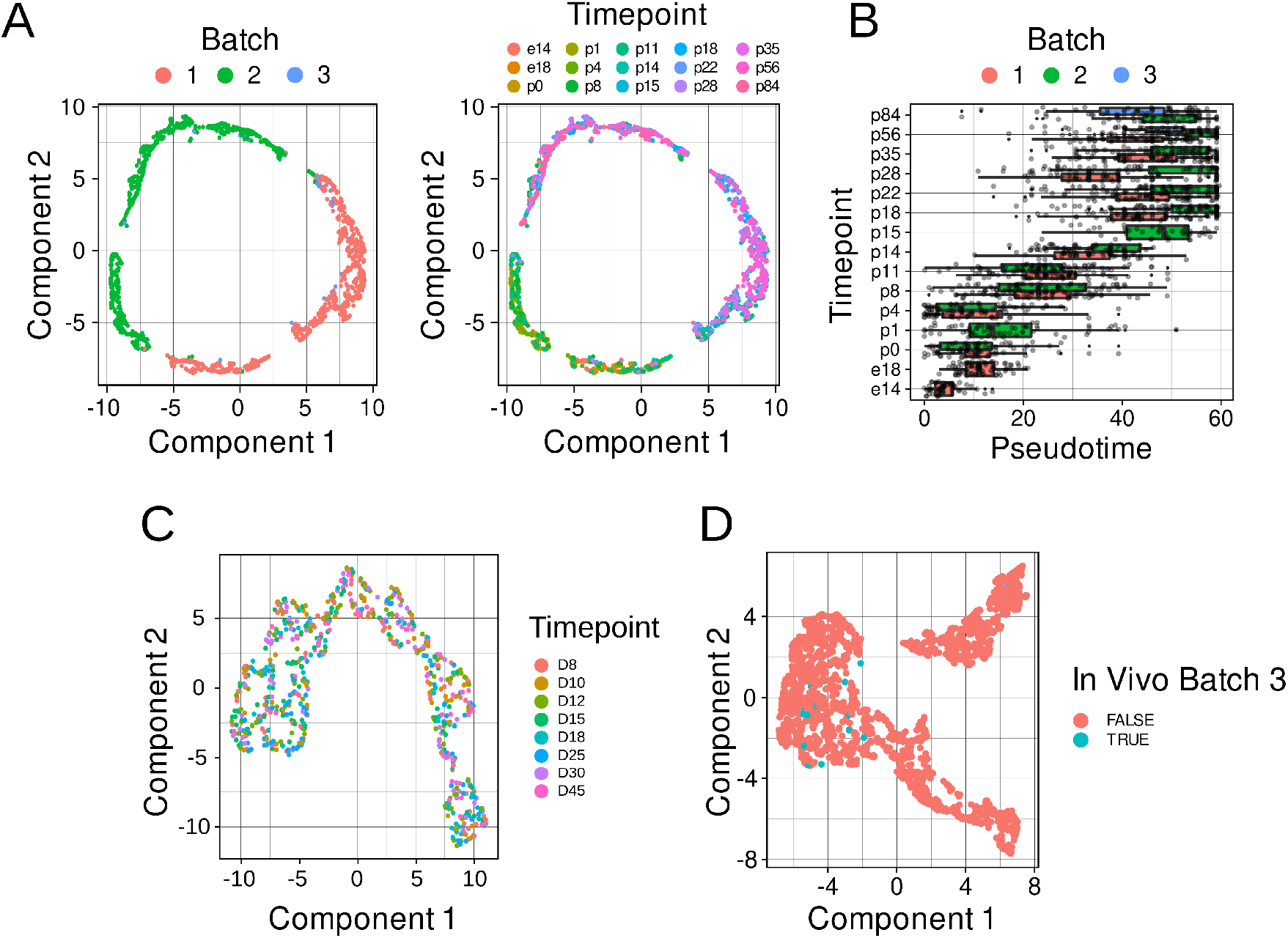
Balanced library design enables correction of scRNA-seq batch effects. **A.** Uncorrected Monocle 3-inferred trajectories of *in vivo* CMs, labeled by batch (left) or timepoint (right). **B.** Pseudotime scores per timepoint for the inferred trajectory, as in **Fig 1C**, further labeled by batch. **C.** Completely unbiased Monocle 3-inferred trajectory for PSC-CMs. **D.** Combined *in vivo* CM and PSC-CM trajectories inferred by Monocle 3, as in **Fig 2F**. Endogenous CMs from Batch 3 (which contained both endogenous and PSC-CMs) are labeled.

